# Axonal dysfunction precedes axon degeneration in a mouse model of progressive parkinsonism

**DOI:** 10.64898/2026.09.29.755486

**Authors:** Anna M. Lipkin, Alejandra Boronat Garcia, Nora L. O’Prey, Renshu Zhang, Zayd M. Khaliq

**Affiliations:** Section on Cellular Neurophysiology, National Institute of Neurological Disorders and Stroke (NIH), Bethesda MD

## Abstract

Axons of substantia nigra dopaminergic neurons “die back” in Parkinson’s disease (PD), exhibiting early degeneration relative to cell bodies, a key feature also observed in PD models. A critical step towards understanding the process of axonal neurodegeneration is to determine the physiological properties of dopaminergic axons that are altered during the early stages of cellular dysfunction. Here, we compared the electrophysiology and calcium dynamics of dopaminergic axons in the MitoPark model of PD with age-matched controls. Our experiments show a pronounced reduction in action potential (AP)-evoked calcium signals in MitoPark axons that emerges well before the onset of dopaminergic fiber loss and motor deficits. In addition, calcium imaging and direct axon recording revealed deficits in AP propagation accompanied by alterations in the axonal AP waveform. Therefore, our experimental findings identify the progression of axon-specific physiological deficits that occur long before neurodegeneration, more precisely defining the period that precedes axonal loss.

## INTRODUCTION

Parkinson’s disease (PD) is a progressive neurodegenerative disease that is caused by the loss of dopamine neurons in the substantia nigra pars compacta (SNc). The selective loss of SNc dopaminergic neurons early in the disease has led to the hypothesis that these neurons have intrinsic properties that make them particularly susceptible to degeneration^1–5^. In particular, SNc dopaminergic neurons fire spontaneously as pacemakers, and they have large, complex, unmyelinated axonal arbors that project into the striatum^6,7^. This combination of pacemaker firing and axonal complexity has been proposed to generate a high baseline energy usage in SNc dopaminergic neurons and make them particularly vulnerable to metabolic stress, which may result from environmental toxins, genetic risk factors, or protein aggregation^2,8,9^. First, action potential (AP) generation drives a substantial fraction of a neuron’s energy expenditure^10–12^. Furthermore, computational modeling has revealed that more complex axonal architecture requires more energy to propagate APs^8^. Finally, experimental evidence shows that blocking AP generation equalizes the energy expenditure between SNc dopaminergic neurons and those in the ventral tegmental area (VTA)^13^. These findings support the idea that the combination of autonomous pacemaking and extensive axonal arborizations contribute to the elevated energetic demands of SNc dopaminergic neurons, and potentially to their selective vulnerability. In particular, given their elevated energetic demands, SNc dopaminergic neurons require high levels of mitochondrial ATP production, which comes at the cost of increased production of reactive oxygen species and mitochondrial oxidant stress^14^. Together, the increased energy expenditure and production required in SNc dopaminergic neurons may ultimately promote mitochondrial dysfunction and eventual neuronal degeneration.

Mitochondria are a common site of dysfunction across a variety of PD etiologies. Toxin-based PD models, including administration of 1-methyl-4-phenyl-1,2,3,6-tetrahydropyridine (MPTP) and rotenone, directly impair mitochondrial function via the inhibition of mitochondrial complex 1, part of the electron transport chain^15–17^. Similarly, genes implicated in familial PD, including PINK1, parkin, and DJ1 lead to impairment across a variety of mitochondrial pathways, while complex 1 activity is reduced in the SNc of patients with idiopathic PD^18–22^. Furthermore, selective disruption of mitochondrial function is sufficient to drive progressive parkinsonian phenotypes in mouse models^23,24^. Thus, mitochondrial dysfunction represents a common feature with potential to compromise the high energy production demands of SNc dopaminergic neurons.

The extensive axons of SNc dopaminergic neurons themselves may be particularly sensitive to mitochondrial dysfunction. In addition to AP propagation, axons use energy to maintain resting membrane potential across a large surface area, traffic cargo across long distances, and coordinate synaptic release machinery^1^. For this reason, axons are thought to be particularly vulnerable in PD: axon loss occurs early and precedes the loss of dopamine cell bodies in both human PD cases and animal models^25–27^. This observation led to the hypothesis of axonal “die-back” in PD, the idea that axon dysfunction and eventual death lead to the degeneration of dopamine cell bodies^26–29^. This hypothesis is supported by observations that mitochondrial motility is impaired in iPSC-derived neurons from familial PD patients, and the number, size and proximity of mitochondria to axonal dopamine release sites are reduced in post-mortem tissue from PD patients^30,31^. However, it remains unclear to what extent mitochondrial dysfunction affects the fundamental electrical functions of the axon: AP generation and propagation.

MitoPark mice carry a dopamine neuron-specific deletion of *Tfam*, the gene encoding mitochondrial transcription factor A^24^. This mutation causes progressive loss of striatal dopaminergic axons, a feature that makes this model convenient to study the physiology of axons before the onset of degeneration. *Tfam* mediates mitochondrial quality control, and its deletion leads to impaired mitochondrial function and energy production ^32^. When this deletion is targeted to neurons that express the dopamine transporter (DAT), mice display progressive neurodegeneration of dopamine neurons in the SNc and progressive motor deficits that begin around 14–20 weeks of age^24,33^. In these mice, there is progressive loss of tyrosine hydroxylase (TH)-positive fibers in the dorsal striatum before the onset of movement deficits, typically between 12–14 weeks of age^24,33^. This fiber loss begins in the most dorsolateral portion of the striatum and continues ventromedially over time^33,34^. At 8 weeks of age, before there is significant reduction in the number of TH+ striatal fibers, MitoPark mice show reductions in evoked striatal dopamine release relative to controls, which may result from changes in axonal excitability but has not been directly tested^35^. Importantly, in the SNc of MitoPark mice, there are no overt changes in somatic firing rate or regularity before 11–15 weeks of age^36^. The well-defined temporal progression of axon loss and motor symptoms in MitoPark mice provides an opportunity to examine dopaminergic axon function during a prodromal stage, where mitochondrial function is impaired but substantial axonal degeneration has not yet occurred. Thus, we focused on testing whether deficits in AP generation propagation precede the loss of dopaminergic axons.

Here, we used *ex vivo* calcium photometry, two-photon calcium imaging and cell-attached electrophysiological recordings of single axons to define the physiological changes that occur before axon degeneration in the MitoPark mouse model of PD. We tested both the excitability of nigrostriatal axons and their ability to propagate signals at three different timepoints preceding axon loss and symptom onset. We found that impairing mitochondrial function leads to a reduction in evoked calcium influx in nigrostriatal axons in MitoPark mice that precedes axon degeneration. We additionally observed that MitoPark axons show a deficit in the generation and propagation of high-frequency bursts of APs as early as 9 weeks of age. These changes in burst propagation were accompanied by differences in AP waveform that suggest that the excitability of nigrostriatal axons is altered before axonal degeneration in the MitoPark mouse model of progressive parkinsonism.

## MATERIALS AND METHODS

### Animals

All animal procedures were approved by the animal care and use committee (ACUC) for National Institute of Neurological Disorders and Stroke (NINDS) at the National Institutes of Health. Mice of both sexes were used for all experiments. Tfam^fl/fl^ mice (Jackson Labs strain #026123) were crossed with DAT^Cre/+^ mice (Jackson Labs strain #006660) to generate Tfam^fl/fl^::DAT^Cre/+^, or MitoPark, mice ^24^. MitoPark mice were used at 6, 9, and 12 weeks of age. Rather than littermate controls, age-matched DAT^Cre/+^ mice were used, as dopamine release has been shown to be altered in hemizygous DAT^Cre/+^ mice relative to wild-types ^37^.

### Viral Injections

Stereotaxic injections were performed on a Stoelting QSI. Mice between 3 and 9 weeks of age were anesthetized with isoflurane and maintained under anesthesia for the duration of the procedure. Anesthetic depth was assessed periodically using the pedal withdrawal reflex and respiration rate. A 50/50 mix of AAV9-CAG-FLEX-tdTomato-WPRE-bGH (Penn Vector Core) and AAV9-pGP-AAV-syn-FLEX-jGCaMP8f-WPRE (Janelia) viruses (0.4 µL/side) were injected bilaterally into the SNc (ML±1.2, AP -3.0, DV 4.0) at 0.1 µL/min via a Hamilton syringe. Following the injection, the syringe was raised at 0.1 to 0.2 mm per minute for 10 minutes before the needle was removed. Animals were allowed to recover from anesthesia on a warmed pad and received ketoprofen on the day of surgery and for two days following. The virus was allowed to express for 3 weeks before photometry and two-photon imaging experiments. For electrophysiology experiments, the tdTomato virus was injected alone and allowed to express for at least 1 week before experiments.

### Ex vivo photometry

Photometry experiments were performed 3 weeks after viral injection on mice of both sexes at 6, 9, or 12 weeks of age. Mice were anesthetized using isoflurane, then the brain was rapidly extracted and placed in cutting solution containing, in mM: 198 glycerol, 2.5 KCl, 1.2 NaH_2_PO_4_, 25 NaHCO_3_, 20 HEPES, 10 glucose, 10 MgCl_2_, 0.5 CaCl_2_, 5 sodium ascorbate, 3 sodium pyruvate, and 2 thiourea. 330 µm thick horizontal slices containing the dorsal striatum were collected using a Leica VT 1200S vibratome. Viral injection site was verified during slicing using the NightSea XITE-RB Fluorescence Flashlight System (Electron Microscopy Sciences). After sectioning, slices were immediately moved to a recovery chamber containing holding solution (containing, in mM: 92 NaCl, 2.5 KCl, 1.2 NaH_2_PO_4_, 30 NaHCO_3_, 20 HEPES, 35 glucose, 2 MgCl_2_, 2 CaCl_2_, 5 sodium ascorbate, 3 sodium pyruvate, and 2 thiourea, 300–310 mOsm) at 32–34°C, where they were allowed to recover for 30 min. Both cutting and holding solutions were bubbled with 5% CO_2_/ 95% O_2_. Slices were then moved to room temperature and allowed to equilibrate for at least 30 min before being moved to the recording chamber. Slices were then transferred to a recording chamber that was continuously perfused with recording solution (containing, in mM: 125 mM NaCl, 25 mM NaHCO_3_, 1.25 mM NaH_2_PO_4_, 3.5 mM KCl, 10 mM glucose, 1 mM MgCl_2_, and 2 mM CaCl_2_, 315-320 mOsm) that was heated to 30-34°C and bubbled with 5% CO_2_/ 95% O_2_. Slices were allowed to equilibrate for at least 15 min in recording solution before imaging. For all experiments, the recording solution contained 30 nM atropine, 300 nM sulpiride, and 300 nM CGP55845 to prevent feedback activity from the network from affecting axon excitability. Nigrostriatal axons receive synaptic-like input from cholinergic neurons which can generate ectopic APs via nicotinic receptors on the axon^38^. To isolate effects on axonal excitability from potential changes in cholinergic input to dopaminergic axons, 1 µM DHβE was included in the recording solution to block nicotinic receptors.

Electrical stimulation was evoked using tungsten bipolar electrodes (75 µm tip separation, MicroProbes) placed at the medial portion of the caudal striatum. Stimulation was evoked using an Isoflex (A.M.P.I) and amplitudes ranged from 0.3 mA to 3 mA. Stimulation was presented as a train of 10 stimuli at 2 Hz, followed by a train of 5 stimuli at 20 Hz, 10 more stimuli at 2 Hz, and finally 5 stimuli at 100 Hz. This protocol was presented 10 times at each imaging location. Traces 2-5 and 7-10 were averaged *post hoc* to increase signal-to-noise ratio. Stimulation-evoked calcium signal was measured using a photodiode (New Focus) mounted above an Olympus BX-51WI microscope. Fluorescence was directed to the photodiode using a DFM1 right-turning filter cube (ThorLabs). Illumination was provided by a DC2200 white LED (Thorlabs) and used with a GFP filter to collect jGCaMP8f fluorescence. The photodiode signal was collected using pClamp 11.1 (Molecular Devices) at 50 kHz and Bessel filtered at 20 kHz. Stimulation electrode location was imaged for each slice using Dodt contrast optics and tdTomato fluorescence using an ORCA-Fusion BT Digital CMOS camera (Hamamatsu).

### Two-photon calcium imaging

Two-photon imaging experiments were performed using a two-photon microscope from Bruker Technologies. A Mai Tai ultrafast Ti:Sapphire laser (Spectra Physics) was tuned to 900 nm for calcium imaging. Fluorescence was collected using a 60X, 1.00 NA objective (Olympus) and a 1.4 NA oil immersion condenser (Olympus). Fluorescence was split into red and green channels using a 575 nm dichroic longpass mirror and pass through 607/45 nm and 525/70 nm barrier filters before detection by photomultiplier tubes (PMTs). H7422PA-40 PMTs (Hamamatsu) were used to capture green fluorescence and multi-alkali R9110 PMTs (Hamamatsu) captured red fluorescence. Red fluorescence was used to identify axonal segments for scanning. Fluorescence data was captured in linescan configuration, where the laser was repeatedly scanned over a segment of axon at ∼450 Hz. Data were collected from 6 repeated presentations of the stimulus pattern and reported as the change in green fluorescence detected divided by the red fluorescence (ΔG/R). Single trial data were smoothed using a 3-point binomial filter in Igor Pro (Wavemetrics) before analysis.

### Ex vivo electrophysiology

Cell-attached recordings were made from nigrostriatal axons in the striatum in mice from both sexes at 9 weeks of age. Mice were anesthetized and the brain was extracted as described above. Sagittal sections (350 µm thick) containing the striatum were collected and allowed to recover as described above. Patch electrodes were pulled from borosilicate glass capillaries (World Precision Instruments) on a horizontal puller (Sutter Instruments) to a resistance of 12-15 MΩ. Pipettes were filled with the ACSF recording solution and wrapped with parafilm to reduce capacitance. A bipolar stimulating electrode was placed into the medial fiber bundle, which was visualized using the expression of tdTomato in nigrostriatal axons. Recordings were made from bouton-like swellings on axons in the striatum. Test pulses of electrical stimulation were delivered to determine if the recorded axon was responsive to the stimulating electrode. Only axons that showed reliable APs following the first stimulus presentation (AP 1 of 25, 100% probability) were included in analysis. Stimulation strength ranged from 0.3 to 5 mA.

Electrophysiological data were acquired with a MultiClamp 700B amplifier and custom routines in pClamp. Data were acquired at 50 kHz and filtered at 3 kHz. To measure APs in cell-attached axonal recordings, the gain was set to 100, and a 1 Hz filter was used to remove any DC offset. Stimulations were presented as a train of 10 stimuli at 2 Hz, 10 stimuli at 20 Hz, 10 stimuli at 2 Hz, and 10 stimuli at 100 Hz. Each axon was recorded for 25 presentations of the same stimulation train with an intertrial interval of 15 seconds.

Electrophysiology data were visually inspected following collection to determine the presence or absence of an AP following each stimulus. The maximum amplitude of the extracellular signal following the stimulus artifact was defined as the AP peak value, and the minimum value following this peak was defined as the AP trough value. These values were used to calculate AP height (AP peak amplitude minus AP trough amplitude) and AP half-width (AP peak time minus AP trough time). To determine AP onset time, traces with 100% AP probability were averaged for each axon and the inflection point of the extracellular voltage trace preceding the AP peak was defined as AP onset.

### Immunohistochemistry

Mice were anesthetized with isoflurane and transcardially perfused with ice-cold phosphate-buffered saline (PBS) followed by 4% PFA in PBS. Brains were removed and post-fixed overnight at 4°C, washed in PBS, and cryoprotected in 30% sucrose until fully equilibrated. Brains were then coronally sectioned, and serial sections were collected at a thickness of 50 µm. Free-floating sections were incubated for 10 min in a 10% Normal Goat Serum solution (Invitrogen, 50062Z), followed by permeabilization for 2 h in PBT containing 0.3% Triton X-100. For striatal sections, consecutive sections were then incubated with one of the following primary antibodies: rabbit anti-mCherry (Proteintech, 26765-1-AP), chicken anti-GFP (Invitrogen, A10262) or sheep anti-tyrosine hydroxylase (TH) (Abcam, ab113), each diluted 1:500 in PBT containing 0.1% Triton X-100. Primary antibody incubation was performed for 3 days at 4°C. Following primary antibody incubation, sections were washed three times for 20 min each in PBS and incubated overnight with the corresponding Alexa Fluor 647-conjugated secondary antibody (Life Technologies; anti-sheep, A21448; anti-rabbit, A32795TR; anti-chicken, A32933TR). Anti-rabbit and anti-chicken secondary antibodies were diluted 1:400, whereas the anti-sheep secondary antibody was diluted at 1:250. Sections were subsequently washed three times for 20 min each in PBS, counterstained with DAPI (1:10,000, Invitrogen, 62248), and washed once more in PBS for 15 min. Sections were mounted for imaging and subsequent quantification. To minimize variability in tissue processing and immunostaining, all animals were processed in parallel using identical staining conditions.

### Imaging and quantification

#### GFP^+^, tdTomato^+^, and TH^+^ fiber quantification

Striatal sections were imaged using a Axioscan 7 Zeiss scanner with a 20x air objective. For each tissue section, a single z-plane centered within the tissue was acquired using the microscope’s automatic middle-z detection based on DAPI signal. Imaging parameters were established using a control section and kept constant across all samples within each experiment. To minimize variability associated with the imaging conditions, all samples were imaged on the same day.

Axonal fibers were quantified in ImageJ. The experimenter was blinded to experimental condition during quantification. For each slice, a boundary was drawn around the striatum using the DAPI signal. Mean fluorescence intensity was quantified within the boundary for each channel using ImageJ’s Measure function. Both hemispheres were quantified for each slice. To account for background fluorescence, mean fluorescence was normalized by dividing by the mean fluorescence intensity of the corpus callosum within the same slice, an area that showed no fluorescent staining.

#### Tissue Clearing and Light Sheet Microscopy

Mice were anesthetized with isoflurane and transcardially perfused with ice-cold phosphate-buffered saline (PBS) followed by 4% PFA in PBS. Brains were removed and post-fixed overnight at 4°C and washed in PBS. For tissue clearing, we followed the LifeCanvas passive clearing protocol. Briefly, samples were first protected by incubation in SHIELD-OFF solution containing SHIELD-EPOXY, SHIELD buffer, and water for 3 days at 4°C. Samples were then transferred to SHIELD-ON solution and incubated for 24 hours at 37°C. For lipid removal, samples were incubated in delipidation buffer at 45°C for 5 days. Following delipidation, samples were incubated in EasyIndex refractive index-matching solution (RI = 1.52; LifeCanvas Technologies) for at least 2 days at 37°C. Samples were first incubated in 50% EasyIndex diluted in distilled water for 24 hours, followed by 100% EasyIndex for an additional 24 hours. Samples were then mounted in EasyIndex Mounting Gel according to the manufacturer’s protocol. After the mounting gel had cured, samples were incubated overnight in EasyIndex at 37°C to optimize imaging quality. Immediately before imaging, samples were washed in EasyIndex Matched Immersion Oil for 10 min at room temperature. Cleared brain hemispheres were imaged using a SmartSPIM light-sheet microscope (LifeCanvas Technologies) with a 3.6 objective. Images were acquired with a z-step size of 4 µm. Virally expressed tdTomato was excited using a 561-nm laser.

#### Chemicals

Sulpiride, atropine, and CGP55845 were dissolved in DMSO. DHβE was dissolved in deionized water. CGP55845 was from Tocris. AAV9-pGP-AAV-syn-FLEX-jGCaMP8f-WPRE was a gift from GENIE Project (Addgene viral prep #162379-AAV9); RRID: Addgene_162379. All other chemicals were from Sigma-Aldrich.

#### Quantification and Statistical Analysis

Analysis was conducted in Igor Pro and Prism 10.1.2 (GraphPad). Two-photon calcium transients were processed in MATLAB (MathWorks) before further analysis in Igor Pro. Unless otherwise stated, data in the text are reported as mean ± standard deviation and displayed with box plots showing median, quartiles, and 10% and 90% tails. Sample sizes, including number of slices, axons, and animals, were chosen based on standards in the field. Specific statistical tests are listed in the text. Bonferroni’s correction for multiple comparisons was applied where appropriate.

## RESULTS

### Nigrostriatal dopaminergic axons in MitoPark mice show age-dependent reductions in baseline calcium and evoked calcium influx

To determine how mitochondrial dysfunction affects nigrostriatal dopaminergic axons in the period before axon degeneration, we performed *ex vivo* calcium imaging experiments in the dorsal striatum of MitoPark (DAT^Cre/+^::Tfam^fl/fl^) and control (DAT^Cre/+^) animals at three prodromal time points (**Fig. 1*A***). To selectively measure calcium signals from dopaminergic axons, animals were injected bilaterally in the SNc with AAV9-pGP-AAV-syn-FLEX-jGCaMP8f-WPRE virus (jGCaMP8f), a genetically-encoded calcium sensor with high signal-to-noise ratio and fast temporal dynamics that make it suitable for detecting calcium influx from single APs. jGCaMP8f was co-injected with a AAV9-CAG-FLEX-tdTomato-WPRE-bGH virus to allow for the morphological identification of striatal dopaminergic axons. To ensure a consistent interval between viral injection and imaging experiments, animals were injected at 3, 6, or 9 weeks of age and *ex vivo* slices were collected at 6, 9, or 12 weeks of age respectively, three weeks after viral injection.

**Figure 1:**
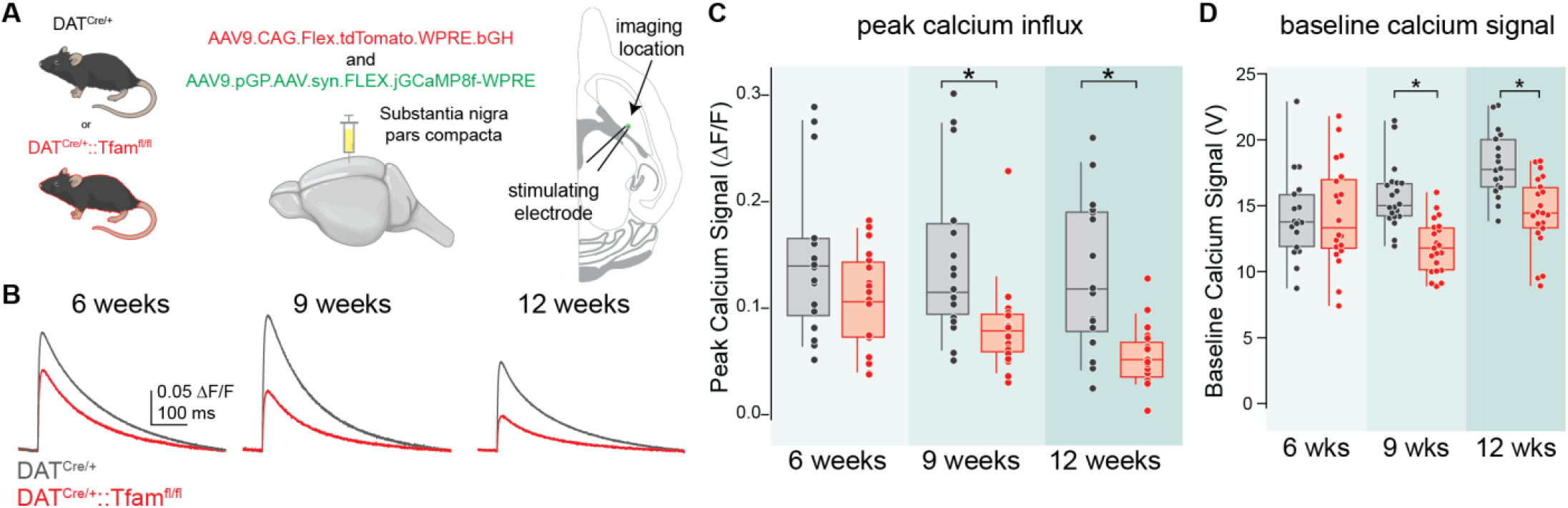
Nigrostriatal axons from MitoPark mice show age-dependent reductions in baseline and evoked calcium influx. ***A***. Experimental setup. Left: the substantia nigra pars compacta of DAT^Cre^/^+^ and DAT^Cre/+^::Tfam^fl/fl^ mice were injected with jGCaMP8f and a virus driving Cre-dependent TdTomato expression to measure calcium influx and label axons originating from the SNc. Right: 3 weeks after injection, calcium influx was evoked in the striatum using a stimulating electrode and measured via a photodiode. Horizontal slices were used to maximize axonal connectivity. ***B.*** Representative traces from DAT^Cre/+^ (black) and DAT^Cre/+^::Tfam^fl/fl^ (red) mice showing axonal calcium influx at three postnatal time points. ***C.*** Summary of peak calcium influx measured across ages for both genotypes reveal an age-dependent reduction in peak evoked calcium influx in MitoPark mice. ***D.*** Summary data of baseline calcium signal. 9-week and 12-week DAT^Cre/+^::Tfam^fl/fl^ slices show reduced baseline calcium signals relative to age-matched controls.

To examine how electrical function changes in MitoPark axons, we generated APs in striatal axons using a bipolar stimulating electrode placed in the dorsal striatum (**Fig. 1*A***). Single electrical pulses were used to evoke activity in striatal dopaminergic axons, and the resulting calcium influx was measured using a photodiode. At 6 weeks of age, there was no significant difference in the peak calcium signal evoked by a single pulse between MitoPark and control mice (**Fig. 1*B–C***, control: 0.144 ± 0.069 ΔF/F; MitoPark: 0.104 ± 0.045 ΔF/F; p = 0.1052, Mann-Whitney). However, at 9 weeks, MitoPark axons showed an approximately 40% reduction in peak calcium amplitude relative to axons from age-matched control animals (**Fig. 1*B–C***, control: 0.141 ± 0.07 ΔF/F; MitoPark: 0.082 ± 0.042 ΔF/F; p = 0.0004, Mann-Whitney). This difference was more pronounced at 12 weeks of age, with the peak calcium amplitude in MitoPark axons reduced by nearly 60% relative to controls (**Fig. 1*B–C***, control = 0.130 ± 0.068 ΔF/F, MitoPark = 0.054 ± 0.027 ΔF/F, p = 0.0002, Mann-Whitney). Overall, the results show that electrically-evoked calcium influx progressively declines in MitoPark axons beginning at 9 weeks of age.

It is possible that mitochondrial dysfunction in MitoPark axons would reduce the calcium buffering capacity of axonal mitochondria and generate a higher level of calcium at baseline, in the absence of AP firing^1^. However, at 9 and 12 weeks, when we measured the amplitude of the photodiode signal before stimulus onset, we instead observed a decrease in baseline calcium photometry signals in MitoPark axons relative to controls. (**Fig. 1*D***, **Supplemental Fig. 1*A***, 6 weeks control: 14.22 ± 3.25 V, 6 weeks MitoPark: 14.31 ± 3.89 V, p = 0.9885, Mann-Whitney; 9 weeks control: 15.67 ± 2.41 V, 9 weeks MitoPark: 11.91 ± 2.01 V, p < 0.0001, Mann-Whitney; 12 weeks control: 18.11 ± 2.47 V, 12 weeks MitoPark: 14.29 ± 2.86 V, p = 0.0002, Mann-Whitney). We also compared the decay of axonal calcium signals between MitoPark and control animals, which would also be affected by alterations in calcium buffering. We found no differences in decay kinetics between MitoPark axons and control axons (6 weeks: p = 0.8523, 9 weeks: p = 0.7051, 12 weeks: p = 0.0734, Mann-Whitney tests).

Together, these results demonstrate an age-dependent reduction in both baseline and electrically-evoked calcium responses in MitoPark nigrostriatal dopaminergic axons that begins at 9 weeks of age.

### Reduced calcium influx precedes detectable loss of nigrostriatal dopaminergic axon fiber density

We next asked if the reduced baseline and evoked calcium signals in MitoPark mice were due to a reduction in the density of dopaminergic axons contributing to the photometry signal. To test whether a reduction in axon fiber density underlies the decreased calcium signal we observed, we quantified the virally-expressed tdTomato fluorescence in striatal slices from MitoPark and control animals at 9 weeks, the first timepoint at which we observed a reduction in baseline and electrically-evoked calcium signals. We observed no difference in the striatal tdTomato fluorescence between MitoPark mice and age-matched controls (**Fig. 2*A–B***, control: 19.57 ± 7.295, MitoPark: 23.59 ± 4.828, p = 0.6286, Mann-Whitney).

**Figure 2:**
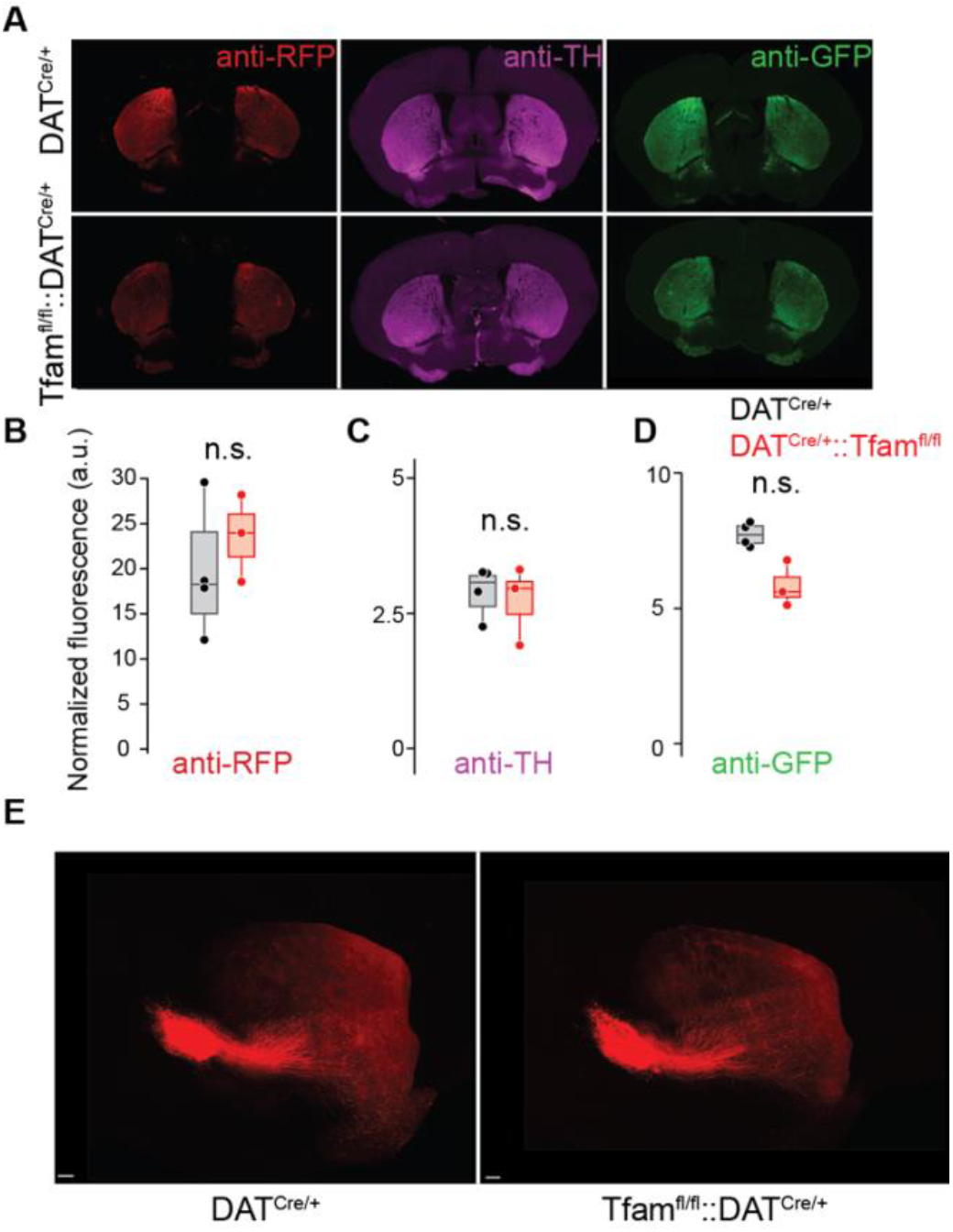
Axon density and dopaminergic identity is not decreased in the striatum of MitoPark mice. ***A.*** Immunohistochemistry of TH and virus expression. Representative images of 9-week DAT^Cre/+^ and DAT^Cre/+^::Tfam^fl/fl^ coronal slices containing the striatum. anti-RFP and anti-GFP were used to measure the expression of the TdTomato and jGCaMP8f viruses, respectively. Tissue was collected at 9 weeks. ***B.*** anti-RFP expression, used to quantify expression of the TdTomato virus, was not different between genotypes. ***C.*** TH expression was comparable between DAT^Cre/+^ and DAT^Cre/+^::Tfam^fl/fl^. ***D.*** anti-GFP expression, used to quantify jGCaMP8f virus expression, was not different in DAT^Cre/+^ and DAT^Cre/+^::Tfam^fl/fl^ striatum. ***E.*** Snapshot of light-sheet microscopy images of tDTomato+ axons in DAT^Cre/+^ (left) and DAT^Cre/+^::Tfam^fl/fl^ (right) brains at 10 weeks. Scale bar, 300 µm

Tyrosine hydroxylase (TH), an enzyme involved in the production of dopamine, is reduced in striatal axons in several PD models, including the MitoPark model, and can be interpreted as either a reduction in dopaminergic striatal axon density or a loss of dopaminergic “identity” ^23,34,39^. Therefore, we next determined whether these axons maintained their dopaminergic identity at this timepoint. To test this, we quantified striatal TH expression using immunohistochemistry and found no significant difference in TH expression between control and MitoPark animals at 9 weeks (**Fig. 2*A, C***, control: 2.914 ± 0.4655, MitoPark: 2.729 ± 0.731, p = 0.057, Mann-Whitney). This finding is consistent with previous studies examining axonal density using TH expression in MitoPark mice at this age^34,40^.

Because differences in viral expression could also contribute to differences in GCaMP fluorescence, we measured viral expression using an anti-GFP antibody. We observed no significant difference between MitoPark and control animals (**Fig. 2*A, D***, control: 7.731 ± 0.434, MitoPark: 5.842 ± 0.854, p = 0.0571, Mann-Whitney). Together, these results indicate that the reduced calcium signal measured with photometry at 9 weeks is not explained by a reduction in dopaminergic axon fiber density or viral expression in MitoPark animals. We therefore next asked if the reduced evoked calcium response we observed in MitoPark axons was accompanied by impaired action potential propagation.

### Propagated signals show substantially reduced amplitudes in MitoPark axons

To determine whether the reduction in evoked calcium influx in MitoPark axons was accompanied by changes in AP propagation, we repeated the single pulse stimulation experiments while measuring calcium influx at increasing distances from the stimulating electrode: at the stimulation site (local site, 0 µm), as above, and two sites far from the stimulating electrode: a proximal site, approximately 150 µm away from the stimulating electrode, and a distal site, about 425 µm away (**Fig. 3*A***). In control axons, peak calcium responses were significantly reduced as the imaging location moved farther from the stimulation site across all ages (**Fig. 3*B–C*,** 0 µm: 0.139 ± 0.068 ΔF/F, 150 µm: 0.095 ± 0.056 ΔF/F, 425 µm: 0.008 ± 0.009 ΔF/F, p < 0.0001, Friedman test with Dunn’s multiple comparisons). A similar pattern was visible in MitoPark axons (**Fig. 3*B–C***). Overall, MitoPark axons showed significantly lower calcium influx than control axons, including as early as 6 weeks (genotype effect, 6 weeks: p = 0.044, 9 weeks: p = 0.0022, 12 weeks: p = 0.0001, Mixed Effects 2-way ANOVAs). To determine if the relationship between distance and calcium influx was different between genotypes, we next tested for an interaction between imaging distance and calcium influx across genotypes for each age. At all three timepoints tested, we found a significant distance x genotype effect (**Fig. 3*C***, distance x genotype effect, 6 weeks: p = 0.0233, 9 weeks: p = 0.0002, 12 weeks: p < 0.0001, Mixed Effects 2-way ANOVAs). In particular, calcium responses measured at the proximal site (∼150 µm away from the stimulating electrode) were significantly lower in MitoPark mice relative to controls at both 9 and 12 weeks (**Fig. 3*C***, 9 weeks: p = 0.0094, 12 weeks: p = 0.0039, Mixed Effects 2-way ANOVA, adjusted p-values reported). Together, these results show that the reduction in axonal calcium influx that we observe with distance differs in MitoPark axons relative to controls at the earliest timepoint assessed.

**Figure 3:**
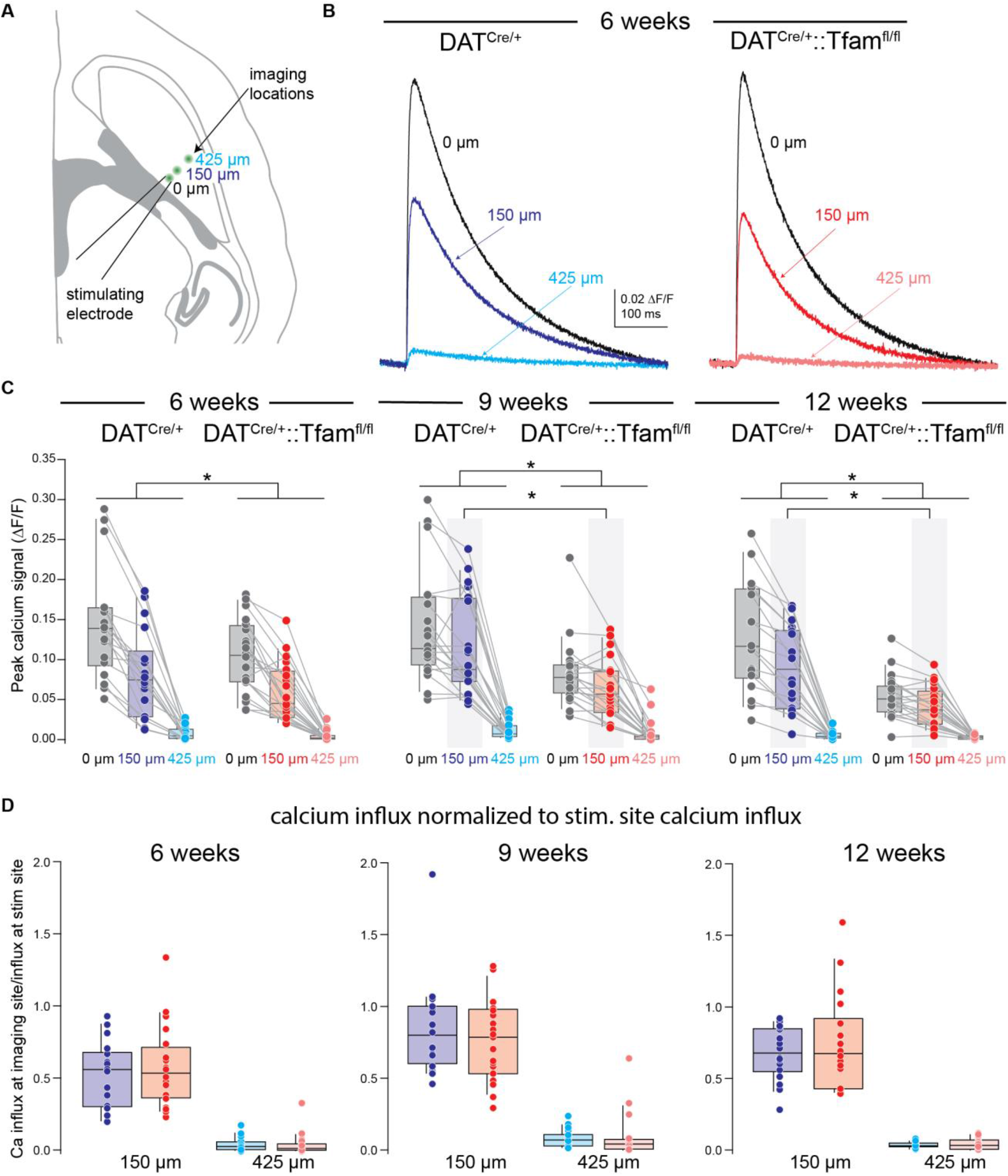
Propagated signals show substantially reduced amplitudes in MitoPark axons. ***A.*** Experimental setup: the photodiode imaging location was moved to measure propagated signals. For between-slice consistency, the spot was moved 100 µm rostral and 100 µm lateral from the stimulating electrode for the 150 µm distance and 300 µm rostral and 300 µm lateral for the 425 µm distance. ***B.*** Representative traces showing peak calcium influx in DAT^Cre^/^+^ (blues) and DAT^Cre/+^::Tfam^fl/fl^ (reds) nigrostriatal axons at three different distances. Note the reduced calcium influx with increased distance from the stimulating electrode. ***C.*** Summary data showing the reduction in calcium influx with distance for each slice at the three timepoints examined. A significant distance by genotype effect was measured at each timepoint. Gray bars show highlight signals collected 150 µm from the stimulating electrode, which were significantly lower in DAT^Cre/+^::Tfam^fl/fl^ slices at 9 and 12 weeks. *D.* Calcium influx at the given imaging location, normalized to the calcium influx at the stimulation site. Each dot represents one slice.

MitoPark axons exhibited reduced calcium influx at the stimulation site at 9 and 12 weeks, and the reduced calcium influx at 150 µm could result from this reduced local response. To test this, we normalized the calcium influx at 150 µm and 425 µm to that measured at 0 µm for each slice. Across ages and genotypes, we found no difference in the normalized calcium influx at 150 µm or 425 µm (**Fig. 3*D***, 150 µm, 6 weeks: p = 0.8285, 9 weeks: 0.4007, 12 weeks: p = 0.4102; 425 µm, 6 weeks: p = 0.4091, 9 weeks: p = 0.1153, 12 weeks: p = 0.5489, Mann-Whitneys). These results show that the difference in peak calcium influx measured at 150 µm and 425 µm in MitoPark axons is largely downstream of reduced calcium responses at the stimulation site at these ages. Thus, it appears that calcium influx is reduced both locally and distally in MitoPark axons after 9 weeks when elicited by single pulses of electrical stimulation. Additionally, MitoPark axons differ from controls in how distance from the stimulation site affects calcium influx, which indicates there may be changes in propagation in MitoPark axons as early as 6 weeks.

### Burst-evoked calcium signals indicate reduced axonal excitability early in MitoPark mice

SNc dopamine neurons fire regularly at 2–4 Hz but can also fire in bursts reaching frequencies up to 50 Hz^7,41,42^. Because burst firing is more likely to reveal differences in the ability of axons to generate APs, we next asked whether repetitive stimulation would reveal excitability deficits in MitoPark axons that may underlie the propagation-related changes we observed with single pulse excitation. To test whether burst firing is affected in MitoPark mice, we presented a train of 5 stimuli at 20 Hz and measured the resulting calcium signal using photometry as described above. At 6 weeks, there was no significant difference in the peak calcium signal between MitoPark mice and age-matched controls (**Fig. 4*A**– B***, control: 0.352 ± 0.166 ΔF/F, MitoPark: 0.246 ± 0.106 ΔF/F, p = 0.0548, Mann-Whitney). At 9 and 12 weeks, however, peak calcium responses to 20 Hz stimulation were reduced in MitoPark mice relative to controls (**Fig. 4*A**– B***, 9-week control: 0.343 ± 0.147 ΔF/F, 9-week MitoPark: 0.208 ± 0.101 ΔF/F, p < 0.0001, Mann-Whitney; 12-week control: 0.328 ± 0.156 ΔF/F, 12-week MitoPark: 0.156 ± 0.075 ΔF/F, p = 0.0002, Mann-Whitney). Although dopaminergic axons do not normally fire at 100 Hz *in vivo*, they are capable of firing as fast as 125 Hz in slice preparations^43^. We hypothesized that 100 Hz stimulation would more effectively reveal whether differences in excitability exist in striatal dopamine axons. We predicted this would result in a more pronounced reduction in calcium influx in MitoPark axons. At both 9 and 12 weeks, peak calcium responses to 100 Hz stimulation were significantly smaller in MitoPark mice than in controls (**Fig. 4*C–D***, 9-week control: 0.352 ± 0.150 ΔF/F, 9-week MitoPark: 0.225 ± 0.106 ΔF/F, p = 0.0009, Mann-Whitney; 12-week control: 0.363 ± 0.165 ΔF/F, 12-week MitoPark: 0.176 ± 0.086 ΔF/F, p = 0.0001, Mann-Whitney). However, contrary to our prediction, increasing the stimulation frequency to 100 Hz did not further increase the magnitude of the reduction, which was similar for 20 and 100 Hz stimulation. Thus, we found that calcium influx in MitoPark axons is reduced during burst firing as well as during single pulse stimulation.

**Figure 4:**
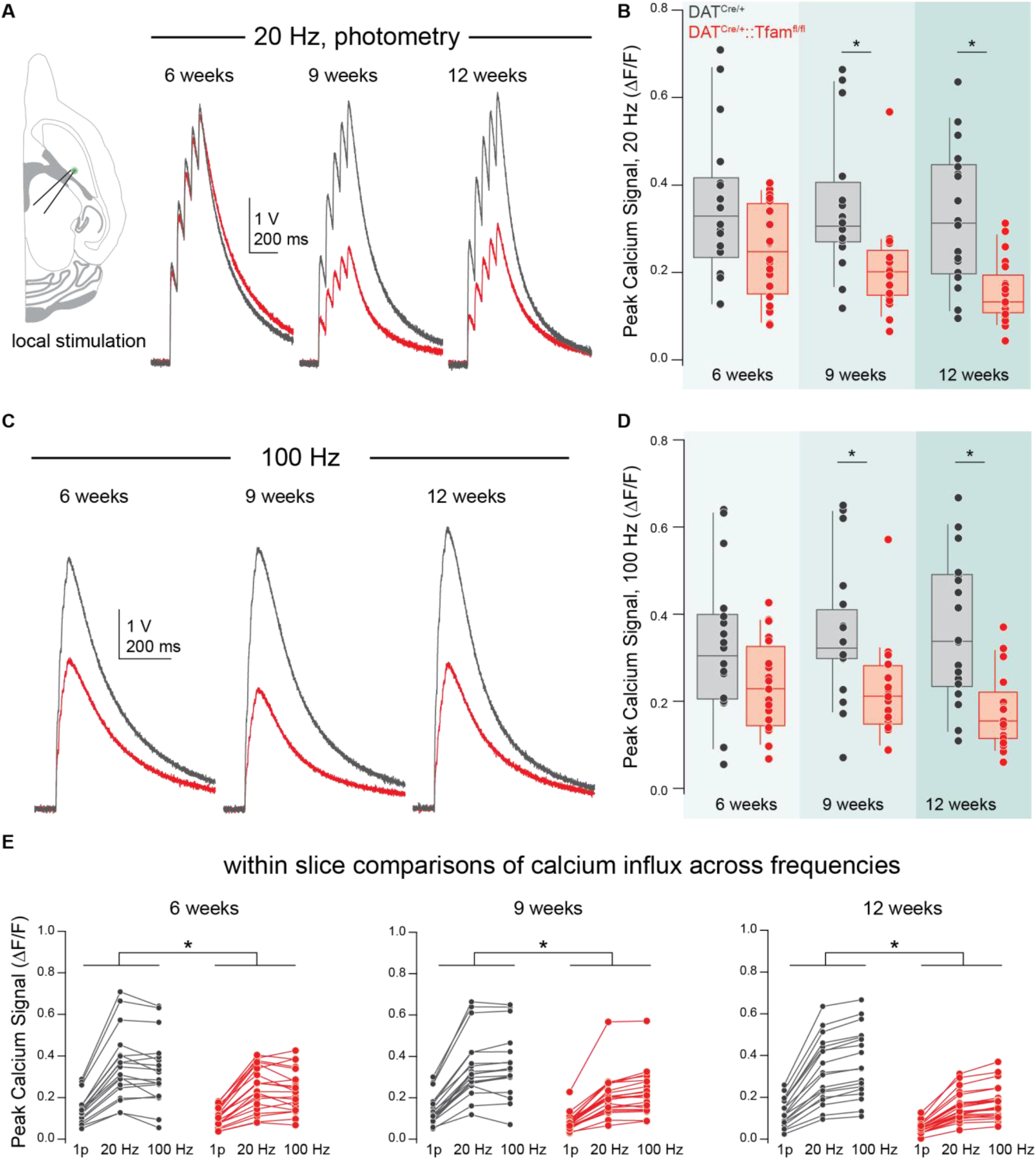
Burst-evoked calcium influx is reduced in MitoPark mice at 9 and 12 weeks of age. ***A.*** Representative traces of calcium influx elicited by 5 pulses of 20 Hz stimulation in DAT^Cre^/^+^ (black) and DAT^Cre/+^::Tfam^fl/fl^ (red) nigrostriatal axons. ***B.*** Summary data showing the peak calcium influx for 20 Hz stimulation across genotype and ages. Calcium influx is significantly reduced in DAT^Cre/+^::Tfam^fl/fl^ axons relative to age-matched controls at 9 and 12 weeks of age. ***C.*** Representative traces of calcium elicited by 5 pulses of 100 Hz stimulation in DAT^Cre^/^+^ (black) and DAT^Cre/+^::Tfam^fl/fl^ (red) nigrostriatal axons. ***D.*** Summary data of 100 Hz shows that calcium influx is reduced in an age-dependent manner in MitoPark axons relative to age-matched controls. ***E.*** The relationship between frequency and calcium influx within each slice, plotted by genotype. There is a significant genotype x frequency effect at all ages tested.

We next sought to determine if the reduced calcium influx we observed in MitoPark axons was the result of changes in axonal excitability. Because we observed reduced single-pulse calcium influx in MitoPark axons at 9 and 12 weeks (**Fig. 1**), it is possible that reductions in burst-associated calcium influx in MitoPark axons are the result of this reduced single-pulse calcium influx rather than excitability changes. Since our 20 Hz and 100 Hz signals were collected in the same slice as our single pulse signals, we were able to compare the amplitudes of burst stimuli to that of a single stimulus for both 20 Hz and 100 Hz bursts. We hypothesized that if excitability were unchanged between MitoPark axons and controls, we should see a similar relationship between single-pulse and burst-evoked calcium influx across genotypes. Conversely, a difference in the relationship between these calcium responses could indicate that excitability deficits underlie the reduced burst-associated calcium influx in MitoPark mice. When we compared the relationship between single-pulse, 20 Hz, and 100 Hz burst responses, we found a significant genotype x frequency effect at all three timepoints tested (**Fig. 4*E***, genotype x frequency effect, 6 weeks: p = 0.0166, 9 weeks: p = 0.0003, 12 weeks: p < 0.0001, Mixed-Effect 2-way ANOVAs). These results indicate that MitoPark axons have excitability deficits that lead to reduced calcium influx during bursts relative to single-pulse stimulation when compared to control axons. The effects occur even at 6 weeks, before a reduction in single-pulse calcium influx is detectable in MitoPark axons.

### Two-photon imaging reveals reduced AP-evoked calcium influx in individual axons in MitoPark mice

Because we saw a pronounced decrease in calcium influx in MitoPark axons in bulk photometry signals, as well as deficits in propagation and excitability, we next turned to two-photon linescan imaging to assess whether this decrease in overall calcium influx was additionally due to a reduction in calcium influx in individual axons, for example, by a reduction in the density or expression of voltage-gated calcium channels. We used the same approach described above to express jGCaMP8f and tdTomato in nigrostriatal axons of MitoPark mice and age-matched controls. We then stimulated nigrostriatal axons with an electrode placed in the dorsal striatum. We performed linescans over tdTomato+ axonal segments <150 µm from the stimulating electrode to quantify calcium influx in single axons (**Fig. 5*A***). To account for the size of the axonal segment, we normalized the jGCaMP8f signals to the red tdTomato signal and reported our results as ΔG/R. Axons with no detectable calcium influx were not analyzed (see Discussion). At 6 and 9 weeks, we saw comparable levels of stimulation-evoked calcium influx for single stimuli in both genotypes (**Fig. 5*B***, 6 weeks control: 0.047 ± 0.034 ΔG/R, 6 weeks MitoPark: 0.06 ± 0.057 ΔG/R, p = 0.6529, Mann-Whitney; 9 weeks control: 0.034 ± 0.019 ΔG/R, 9 weeks MitoPark: 0.022 ± 0.01 ΔG/R, p = 0.1996, Mann-Whitney). However, at 12 weeks, we found a 50% reduction in peak calcium influx in individual MitoPark axons relative to controls (**Fig. 5*B***, 12 weeks control: 0.031 ± 0.026 ΔG/R, 12 weeks MitoPark: 0.015 ± 0.007 ΔG/R, p = 0.016, Mann-Whitney). Unlike our photometry data, we found no change in baseline (pre-stimulus) calcium levels when imaging single axons at any timepoint (**Fig. 5*C***). It should be noted that in our two-photon imaging experiments, we specifically selected for axons that showed calcium influx triggered by electrical stimulation. By contrast, our photometry data collected calcium influx from all axons, regardless of their responsiveness. This discrepancy between our photometry and two-photon data could reflect that non-responsive axons are the primary drivers of low baseline calcium levels in MitoPark axons.

**Figure 5:**
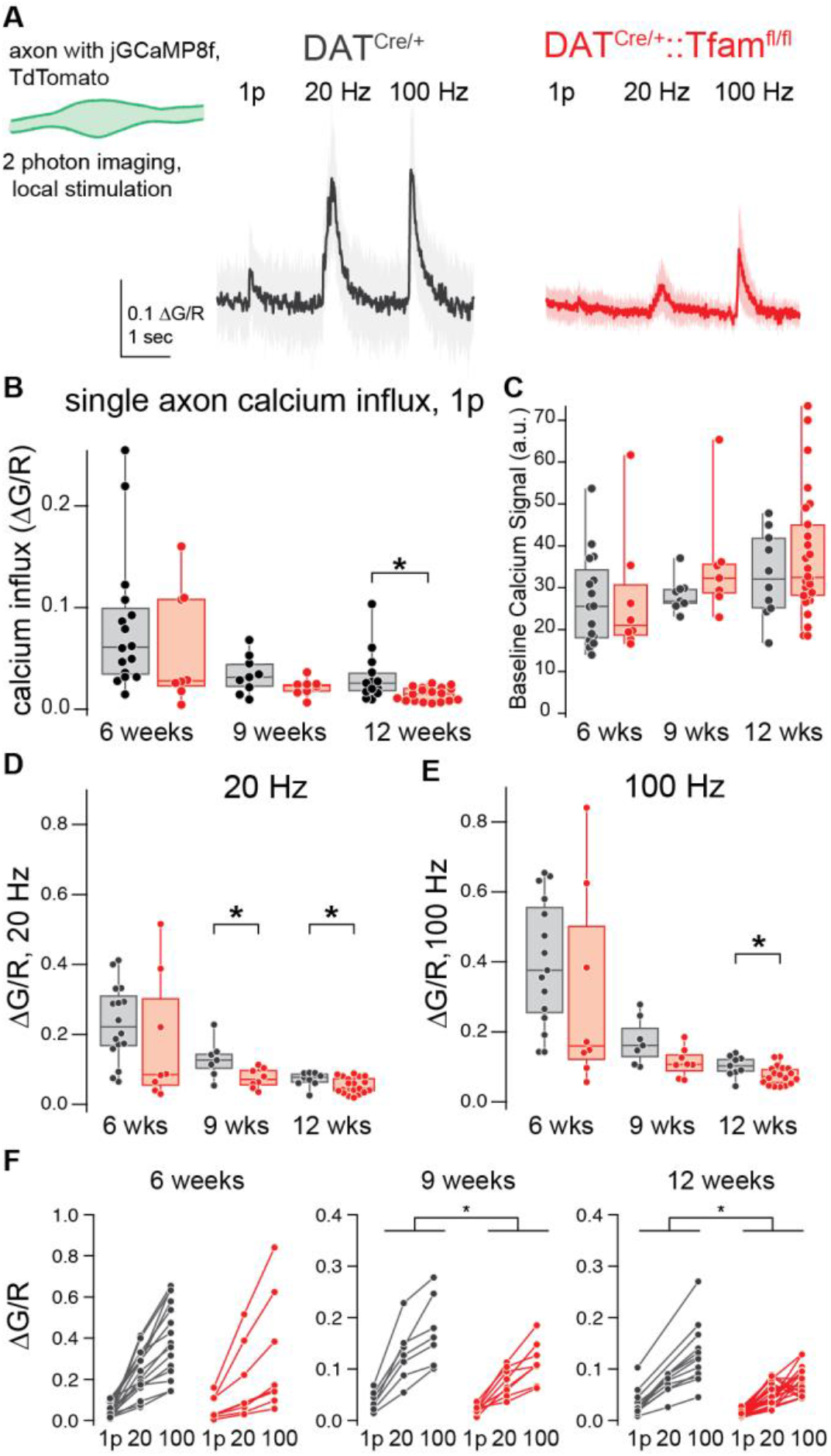
Calcium influx and excitability are reduced at the single axon level in MitoPark axons. ***A.*** Example signals measured by two photon linescan imaging of single axons during high-frequency stimulation. Each trace is the average of six scans per axon. 12-week-old control animal. ***B.*** Calcium influx measured in single axons after a single pulse of electrical stimulation. ***C.*** Baseline calcium levels were quantified as green fluorescence immediately preceding the electrical stimulus. ***D.*** Summary data of the peak calcium signal measured during 20 Hz stimulation across genotypes and timepoints. A significant decrease in calcium was visible in 9- and 12-week DAT^Cre/+^::Tfam^fl/fl^ axons. ***E.*** As in B, but for 100 Hz stimulation. Here, 12-week DAT^Cre/+^::Tfam^fl/fl^ axons showed reduced per-axon calcium influx relative to DAT^Cre/+^controls. *F.* The relationship between calcium influx and stimulation frequency for MitoPark axons differs from control axons at 9 and 12 weeks of age.

We next asked whether the decreases in calcium influx during 20 Hz and 100 Hz stimulation that we observed in MitoPark axons (**Fig. 4**) were also detectable at the single-axon level. First, we confirmed that our individual axons did not show saturating responses to calcium at 100 Hz (**Supplemental Fig. 2**). When we evoked electrical activity with 5 pulses at 20 Hz, we observed a reduction in peak calcium influx at both 9 and 12 weeks in MitoPark axons relative to control axons (**Fig. 5*D***, 9-week controls: 0.129 ± 0.055 ΔG/R, 9-week MitoPark: 0.075 ± 0.028 ΔG/R, p = 0.0289, Mann-Whitney; 12-week control: 0.072 ± 0.021 ΔG/R, 12-week MitoPark: 0.054 ± 0.023 ΔG/R, p = 0.0421, Mann-Whitney). Similarly, stimulation of 5 pulses at 100 Hz resulted in reduced calcium influx in MitoPark axons at 12 weeks of age (**Fig. 5*E***, 12-week control: 0.129 ± 0.059 ΔG/R, 12-week MitoPark: 0.077 ± 0.027 ΔG/R, p = 0.005, Mann-Whitney).

We next asked whether the relationship between the single-pulse calcium signal and burst calcium signal differed between control axons and MitoPark axons, but now at the level of individual axons. Examining the interaction between genotype and frequency, there was a significant effect at 9 and 12 weeks (**Fig. 5*F***, genotype x frequency effect, 6 weeks: p = 0.2677, 9 weeks: p = 0.0311, 12 weeks: p = 0.0054). These results are consistent with our photometry data and suggest that both excitability is impaired in MitoPark axons before axons start to degenerate. Furthermore, these data suggest that the reduction in bulk calcium signals recorded with photometry results in part from this reduction in per-axon calcium influx, which may reflect changes in axonal calcium channel density.

### Burst-evoked calcium influx is reduced in a distance-dependent manner in MitoPark axons

The reduction in excitability that we observed above in MitoPark axons may become more pronounced as APs propagate over long distances in the axon. To assess the ability of MitoPark axons to propagate high-frequency signals, we compared our calcium photometry signals measured at the stimulating electrode to those collected ∼150 and∼ 425 µm away. Axons were stimulated with 5 pulses at 20 Hz and 100 Hz, and overall calcium influx as well as the calcium influx for each peak in the train of 5 stimuli was measured (**Supplemental Fig. 3*A***). To control for the number of stimulated axons in each imaging location, the amplitude of peaks 2 through 5 were normalized to peak 1 (**Supplemental Fig. 3*A***). We reasoned that if all stimulated APs propagated reliably to the more distant imaging sites, we should see a similar pattern of calcium influx across the stimulation train regardless of imaging site. Therefore, we hypothesized that a decrease in the normalized calcium influx for later peaks likely represents a failure, either complete or partial) of some APs to propagate to the distal imaging site. Indeed, we saw a significant effect of imaging location and peak number on normalized calcium influx across all groups (REML mixed-effects model, location x peak number: 6 weeks, p < 0.0001; 9 weeks, p < 0.0001; 12 weeks, p < 0.0001). While normalized calcium signals changed with propagation distance for both genotypes, we next examined whether these reductions differed between MitoPark and control axons. For 20 Hz stimulation, there was no difference between MitoPark and age-matched control axons in the relationship between distance and peak number (**Supplemental Fig. 3*A-B***, REML mixed-effects model, genotype x location x peak number: 6 weeks, p = 0.942; 9 weeks, p = 0.4272; 12 weeks, p = 0.3068). The same was true for 100 Hz stimulation: MitoPark axons and control axons showed comparable distance-dependent decreases in calcium influx later in the stimulation train (**Supplemental Fig. 3*D***, REML mixed-effects model, genotype x location x peak number: 6 weeks, p = 0.9563; 9 weeks, p = 0.9817; 12 weeks, p = 0.0819). These results suggest that there may be propagation failures or changes in AP shape during propagation that result in lower per-stimulus calcium influx later in a train of stimuli in nigrostriatal axons in control mice. However, we saw no difference between genotypes in the magnitude of this distance-dependent reduction in per-stimulus calcium influx.

Although bulk calcium signals are convenient due to their high signal-to-noise ratio, the asynchronous timing of action potential firing across a group of fibers complicates the interpretation of propagated calcium signals. To overcome this temporal blurring, we next tested if deficits in propagation could be detected in individual axons. Using two-photon calcium imaging, we measured the calcium influx resulting from a single stimulation pulse and five pulses at 20 Hz and 100 Hz at a variety of distances from the stimulating electrode (**Fig. 6*A***). For 9-week-old animals, we saw no effect of distance on calcium influx for single-pulse and 20 Hz stimulation in MitoPark axons relative to controls (**Fig. 6*B–C***, control single pulse slope: -3.7 x 10^-5^ ± 330 x 10^-5^ (ΔG/R)/µm, MitoPark single pulse slope: 12 x 10^-5^ ± 2.3 x 10^-5^ (ΔG/R)/µm, p = 0.627, unpaired t-test; control 20 Hz slope: -8.7 x 10^-5^ ± 10 x 10^-5^ (ΔG/R)/µm, MitoPark 20 Hz slope: -9.1 x 10^-5^ ± 9.3 x 10^-5^ (ΔG/R)/µm, p = 0.8309, unpaired t-test). However, with 100 Hz stimulation, we saw that calcium influx decreased with distance to a greater extent in MitoPark relative to control axons (**Fig. 6*D***, control 100 Hz slope: -1.9 x 10^-5^ ± 13 x 10^-5^ (ΔG/R)/µm, MitoPark 100 Hz slope: -13 x 10^-5^ ± 13 x 10^-5^ (ΔG/R)/µm, p = 0.0293, unpaired t-test). At 12 weeks, this difference emerged at even lower frequencies, with both 20 Hz and 100 Hz stimulation resulting in lower calcium influx when measured farther from the stimulating electrode (**Fig. 6 *E–G***, control 20 Hz: 4.5 x 10^-5^ ± 11 x 10^-5^ (ΔG/R)/µm, MitoPark 20 Hz: -7.3 x 10^-5^ ± 3.5 x 10^-5^ (ΔG/R)/µm, p < 0.0001, unpaired t-test; control 100 Hz: 5.6 x 10^-5^ ± 11 x 10^-5^ (ΔG/R)/µm, MitoPark 100 Hz: -9.6 x 10^-5^ ± 4.4 x 10^-5^ (ΔG/R)/µm, p < 0.0001, unpaired t-test). This reduction at more distal locations was not explained by a distance-dependent decrease in the single stimulus-evoked calcium influx. Measuring the ratio between burst-associated calcium influx (5p) and single stimulus influx (5p/1p ratio) over distance, we saw a reduced 5p/1p ratio with distance in MitoPark but not control axons (**Supplemental Fig. 4**). These data suggest that calcium influx is strongly reduced following propagation in MitoPark mice but not controls. This reduction in calcium influx with distance could be explained by changes in AP shape and/or probability. To further examine these possible mechanisms, we next turned to measurement of APs directly via electrophysiological recordings of axons.

**Figure 6:**
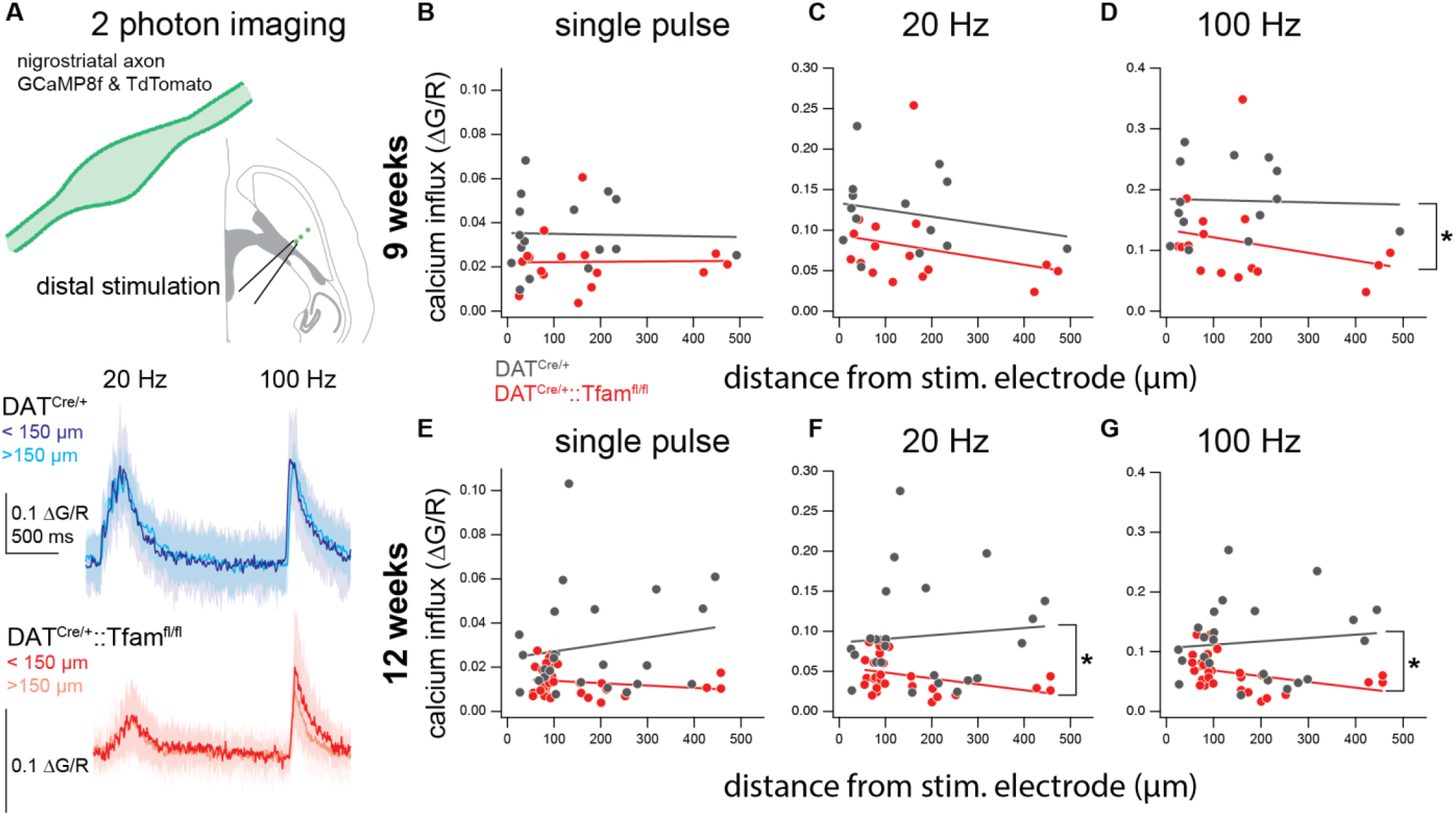
Calcium influx decreases in a frequency- and distance-dependent manner in MitoPark axons. ***A.*** Top: Schematic of experimental approach. Bottom, representative traces of DAT^Cre/+^ (blues) and DAT^Cre/+^::Tfam^fl/fl^ (reds) nigrostriatal axons at two distances from the stimulating electrode. For each genotype, data shown were collected from the same slice (12 weeks of age). ***B.*** Relationship between peak calcium influx and distance from the stimulating electrode for single pulses of stimulation in 9-week animals of both genotypes. The line of best fit was calculated for each genotype using simple linear regression. ***C.*** As in B, but for five pulses of stimulation at 20 Hz. ***D.*** The relationship between peak calcium influx and distance for five pulses at 100 Hz in 9-week animals. There was an inverse relationship between distance and calcium influx in MitoPark but not control axons (unpaired t-test of slopes). ***E.*** As in B, the relationship between peak calcium influx and distance in 12-week-old animals of both genotypes. ***F.*** There was an inverse relationship between distance and calcium influx for 20 Hz pulses in MitoPark axons at 12 weeks. ***G.*** As in F, but for 100 Hz bursts of stimulation.

### Propagation failures occur at lower-frequency firing rates in MitoPark mice relative to age-matched controls

While two-photon calcium imaging offers the benefit of single-axon resolution, the link between calcium influx and AP firing is correlational. To more directly test the ability of striatal axons to propagate bursts at high frequencies, we performed cell-attached recordings from axonal varicosities in MitoPark mice and control animals at 9 weeks of age, the first time point at which we observed changes in calcium influx in our photometry data. To target dopaminergic axons for recording, we injected AAV9-CAG-FLEX-tdTomato-WPRE-bGH virus into the SNc of MitoPark and control animals and prepared sagittal slices to maintain axonal connectivity with the medial fiber bundle, visible as a band of tDTomato+ fibers. The stimulating electrode was placed in the medial fiber bundle to maximize the ability to drive activity across multiple axons (**Fig. 7*A***). tdTomato-positive axons could then be traced into the striatum, where axonal varicosities were visible. Loose-seal cell-attached recordings were made from these axonal varicosities while axons in the medial fiber bundle were stimulated with 2 Hz, 20 Hz, and 100 Hz trains (**Fig. 7*B***). Cell-attached AP waveforms were visible in a subset of axonal varicosities, ranging in distance from 640 to 2270 µm from the stimulating electrode. First, we assessed the probability of AP propagation from the stimulation site to the site of recording by quantifying the number of stimulations per axon that produced a detectable propagated AP. At 2 and 20 Hz, all eight axons we recorded in control animals showed 100% reliable APs across repeated presentations of the stimulation protocol (**Fig. 7*C***). By contrast, two of the eight MitoPark axons we recorded showed failures during the 20 Hz stimulus presentation (**Fig. 7*C***). Axons that demonstrated AP propagation failures showed more failures at the end of the stimulation train but maintained an AP probability rate above 85% in both cases.

**Figure 7:**
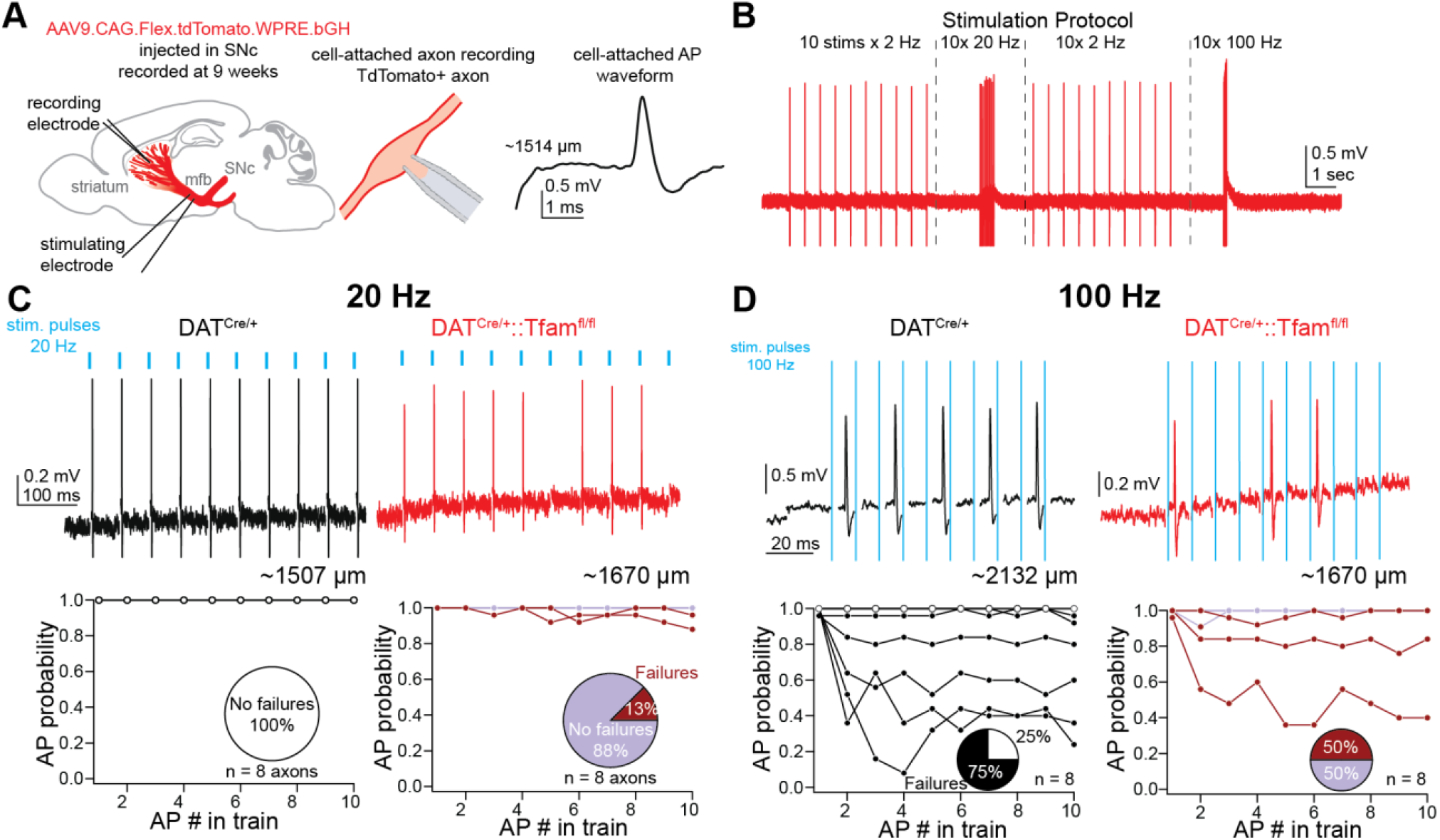
Frequency-dependent propagation failures occur at lower-frequency firing rates in MitoPark mice. ***A.*** Experimental set-up. DAT^Cre/+^ and DAT^Cre/+^::Tfam^fl/fl^ mice were injected with a Cre-dependent TdTomato virus in the substantia nigra pars compacta (SNc). In sagittal slices containing striatum, a stimulating electrode was placed in the medial fiber bundle (mfb) to drive axonal AP firing. Loose-seal cell-attached recordings were performed on axonal varicosities. Right, example cell-attached waveform recorded from an axonal varicosity ∼1514 µm from the stimulating electrode. ***B.*** Diagram of the stimulation protocol used to measure frequency-dependent AP propagation. The stimulation protocol was presented 25 times to each recorded axon. ***C.*** 20 Hz stimulation of nigrostriatal axons generates propagation failures in MitoPark but not control slices. Top: example recordings of DAT^Cre/+^ (black) and DAT^Cre/+^::Tfam^fl/fl^ (red) axons. Each example shows a single trial of 20 Hz stimulation. Blue ticks indicate the timing of electrical stimulation pulses. Bottom, AP probability plotted against AP number in a train of 10 stimuli. For MitoPark axons (pinks), there is a lower probability of AP propagation later in the AP train. Inset: pie charts show the proportion of axons that show any AP failures. ***D.*** 100 Hz stimulation produces propagation failures in both control and MitoPark axons. AP probability plotted against AP number again shows a decrease in AP probability later in a train of stimuli. Inset pie charts show that a higher fraction of control axons shows AP failures relative to MitoPark axons.

While dopamine cell bodies can fire up to 50 Hz in short bouts, they quickly enter depolarization block due to the inactivation of sodium channels^44,45^. Shorter inter-spike intervals minimize the time for sodium channels to recover in between APs, leading to progressively shorter APs and an eventual failure to generate further firing. Thus, we hypothesized that nigrostriatal axons should demonstrate higher AP failure rates at faster stimulation frequencies. At 100 Hz, we saw an increase in the proportion of MitoPark axons that showed AP failures (50%, **Fig. 7*D***) and a decrease in AP probability for those axons that showed failures (average: 83 ± 21.5% AP probability, 4 axons). We also saw a similarly high proportion of axons with AP failures in the control group (**Fig. 7*D***, 6/8 axons with failures, average: 72 ± 27% AP probability). Together, these results suggest that nigrostriatal axons have robust propagation fidelity at 20 Hz in control conditions but produce AP propagation failures at higher frequencies and in the context of mitochondrial dysfunction.

### Cell-attached AP waveforms change with repetitive stimulation in MitoPark and control axons

While we saw a slight elevation in propagation failure probability in our direct recordings of MitoPark axons, these changes were not large enough to explain the frequency-and distance-dependent decreases in calcium influx we observed in these axons using two-photon imaging. Another possible contribution to decreased calcium influx is changes in the AP waveform, where small changes in AP width or height could lead to differences in voltage-gated calcium channel activation and large changes in calcium influx. To test this and to probe the mechanisms underlying the excitability changes we observed in MitoPark and control axons, we next quantified changes in the AP waveform in our cell-attached recordings. To measure how AP waveforms change over the course of a high-frequency burst, we averaged all the trials within each axonal recording in which all APs propagated successfully from the stimulation site to the recording electrode site. We then compared the average waveform of the first and final (10^th^) AP and quantified several features including the delay between the evoked stimulus onset and the maximum amplitude of the AP waveform, the time between AP maximum and AP minimum (roughly equivalent to intracellularly-recorded AP half-width), and the amplitude of the AP, which was measured both from AP onset to maximum as well as from maximum to minimum, equivalent to the upstroke and downstroke velocity of the intracellularly-recorded AP respectively (**Fig. 8*A***). At 20 Hz, we found a pronounced delay in time between the stimulus and peak between the first AP and the tenth within individual MitoPark axons but not controls (**Fig. 8*A–B***, median difference (AP 10 – AP 1): control = 0.04 ms, p > 0.999; MitoPark = 0.08 ms, p = 0.0039, Wilcoxon matched-pairs signed rank test). We also observed an increase of the time from onset to peak in our cell-attached APs in MitoPark axons, which we interpreted to be a slowing of the upstroke velocity in an activity-dependent manner (**Fig. 8*B***, control AP 1: 0.464 ± 0.24 ms, control AP 10: 0.46 ± 0.27 ms, p = 0.992, Wilcoxon test; MitoPark AP 1: 0.393 ± 0.077, MitoPark AP 10: 0.427 ± 0.075, p = 0.0039, Wilcoxon test). We saw no evidence of AP broadening over the course of the 20 Hz stimulation train in either genotype (**Supplemental Figure 5**). At 100 Hz, the increase in delay to peak from AP 1 to AP 10 we observed in MitoPark axons was more pronounced (**Fig. 8*C-D***, median difference (AP 10 – AP 1) control = 0.14 ms, p = 0.3828; MitoPark = 0.22 ms, p = 0.0017, Wilcoxon). Additionally, we saw a decrease in AP height between AP 10 and AP 1, measured from AP onset to AP peak, in both genotypes (**Fig. 8*D***, median difference (AP 10 – AP 1) control =-0.03912 mV, p = 0.0312; MitoPark = -0.04095 mV, p = 0.0391, Wilcoxon). This decrease in onset-to-peak AP height is strongly suggestive of a decrease in sodium channel availability in both control and MitoPark axons across the duration of a 100 Hz stimulation train and may contribute to the high AP failure rate we observed in both genotypes at this high frequency. In addition to the reduction in onset-to-peak height, we also observed a decrease in peak-to-trough height in MitoPark axons but not control axons (**Fig. 8*D***, median difference (AP 10 – AP 1) control = -0.01752 mV, p = 0.0781; MitoPark = -0.05943 mV, p = 0.0078, Wilcoxon). This suggests a reduction in the speed of AP repolarization in MitoPark axons, which may result from changes in potassium channel function in MitoPark axons. The concomitant slowing of AP upstroke and repolarization kinetics without a change in AP half-width suggest that AP height may be decreased in MitoPark axons, consistent with our more pronounced effects of distance and frequency on calcium influx than propagation fidelity in these axons. However, differences in AP height could not be assessed from our cell-attached recordings. Overall, these changes in AP waveform, combined with the changes we observed in calcium influx and propagation, suggest that the earliest functional changes in MitoPark animals are deficits in axonal excitability, and these changes precede axonal degeneration.

**Figure 8:**
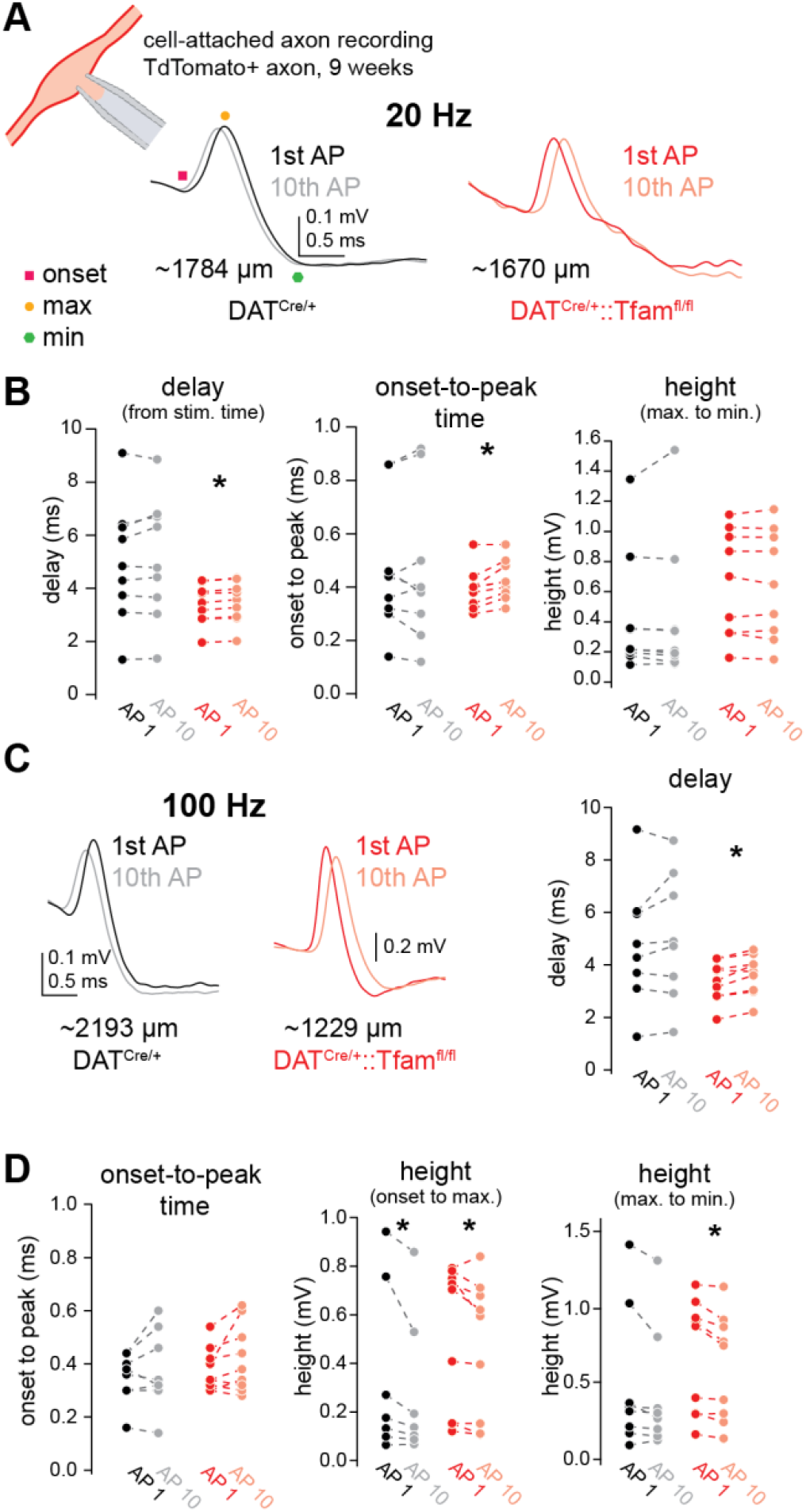
Cell-attached AP waveforms change with high-frequency firing in MitoPark and control axons. ***A.*** Cell-attached axon recordings were performed from TdTomato-positive axonal varicosities in DAT^Cre/+^ and DAT^Cre/+^::Tfam^fl/fl^ mice. For each axon, the cell-attached voltage trace was averaged across trial presentations that showed 100% AP propagation probability. Using the resulting averaged response, it was possible to compare the waveform of the 1^st^ and 10^th^ AP in the 20 Hz train of APs. Representative averages are shown for one axon of each genotype. Recordings were performed at 9 weeks of age. Legend at right indicates onset, maximum, and minimum value of AP waveform. ***B.*** Left: Summary of the difference in delay to peak between AP 10 and AP 1. The delay from stimulus onset to AP peak was greater for the 10^th^ AP relative to the 1^st^ AP in MitoPark but not control axons at 20 Hz. Right: There was no change in AP height, measured from peak to trough of the cell-attached waveform, between APs 1 and 10 for either genotype. ***C.*** Representative averages of the first and 10^th^ AP in a 100 Hz train for DAT^Cre/+^ (black) and DAT^Cre/+^::Tfam^fl/fl^ (red) axons. ***D.*** Left: summary of differences in delay to peak for 100 Hz stimulations. MitoPark axons show an increase in delay to peak between AP1 and AP 10 while control axons show no change. Center: AP height, measured from AP onset to AP peak, changes across the 100 Hz train in both MitoPark and control axons. Right: AP height, measured from AP peak to AP trough, reduces between AP1 and AP10 in MitoPark axons but not control axons.

## DISCUSSION

Here, we show that in a mouse model of progressive parkinsonism, axonal dysfunction precedes axonal degeneration. Notably, we observe a pronounced reduction in electrically-evoked calcium influx in dopaminergic axons as early as 9 weeks of age. This reduction in calcium influx is accompanied by reduced excitability of these axons, as evidenced by lower calcium influx during burst stimulation. Additionally, we observe a deficit in axonal propagation in MitoPark mice using both calcium imaging and electrophysiological recordings. We observe change in the fidelity of AP propagation at high frequencies and changes in the waveform of propagated APs. Together, these effects suggest that alterations in calcium signaling and axonal excitability are among the earliest functional changes in the MitoPark mouse model of progressive parkinsonism.

Early axonal dysfunction is often cited as a key driver of neurodegeneration in PD. However, nearly all evidence of nigrostriatal axonal dysfunction in PD mouse models comes from observations of reduced striatal dopamine release^23,46–48^. Reduced dopamine release could itself result from impaired dopamine synthesis, reuptake, or vesicle filling rather than dysfunction of the axon *per se*. In this study, we assessed axonal function directly, examining axonal electrical activity and calcium influx for the first time in a mouse model of PD. Dopamine release is dependent on both APs and calcium influx, and we have shown that, upstream of any dysfunction in dopamine synthesis or packaging, axonal function is progressively impaired in MitoPark mice before axon loss begins^49–51^. These changes in axonal excitability could contribute to the reduced dopamine release described in 8-week-old MitoPark mice previously^35^. It remains to be determined if the calcium reductions we observed are visible in other mouse models of PD, particularly in other models that are progressive in nature and allow for extended prodromal monitoring of axonal function^23,39,46^. The approach we have established here could be adapted for use in other PD models to assess when and how axonal function is changed and whether alterations of axonal electrical function are a main driver of established dopamine release deficits in these models.

The electrophysiological properties of MitoPark SNc neurons have been well-characterized at the soma^36^. MitoPark SNc neurons showed age-dependent changes in passive membrane properties as well as increasingly irregular firing rates, though largely at ages later than those used in the present study. However, specific decreases in ionic currents were detected as early as 6–10 weeks in MitoPark neurons, specifically the current mediated by small-conductance calcium-sensitive potassium (SK) channels, which contributes to the regularity of AP firing in these neurons^36,52^. In the same study, the authors quantified changes in mRNA expression of these channels and found no change in SK channel expression even as late as 18–22 weeks. One explanation for this discrepancy between recorded currents and mRNA expression could be a reduction in calcium influx that leads to reduced activation of SK channels. Our axonal calcium imaging supports this idea, and the reduction in the AP repolarization speed we observed in our cell-attached axon recordings suggests that SK channels may similarly be activated less in MitoPark axons relative to controls because of their reduced calcium influx. The same study that quantified mRNA expression of SK channels also measured the expression of two calcium channels, *CACNA1C* and *CACNA1D*, which encode Ca_V_1.2 and Ca_V_1.3 channels, respectively. MitoPark animals showed an increase in SNc expression of Ca_V_1.2 expression at 12 weeks, which seems to contradict the decrease in axonal calcium influx we observed in MitoPark axons at this timepoint. However, it is possible that this upregulation in CaV1.2 mRNA is compensatory, an attempt to counteract the apparent reduction in calcium channels in MitoPark axons. Conversely, in the MCI-Park mouse, there is a decrease in Ca_V_1.3 mRNA at P30. This mouse is generated via deletion of *Ndufs2* from dopamine neurons specifically and shows a similar progressive parkinsonian phenotype and loss of striatal dopamine release as the MitoPark mouse, though the time course of neurodegeneration is faster than MitoPark mice and axons do not seem to degenerate^23^. In these mice, the decrease in Ca_V_1.3 mRNA is accompanied by a reduction in action-potential evoked calcium influx, measured in the proximal dendrite. However, the consequences of this mitochondrial dysfunction on axonal calcium remain to be explored.

Our result showing reductions in electrically-evoked calcium influx in MitoPark axons is somewhat unexpected, as excess calcium influx has been associated with the selective vulnerability of dopamine neurons in PD^2,39,53,54^. In particular, pacemaking currents mediated by low-threshold Ca_V_1.3 channels contribute to high intracellular calcium in SNc but not VTA neurons^55,56^. Additionally, blocking calcium influx via isradipine, a specific blocker of Ca_V_1 channels, is protective against both mitochondrial disruptors and alpha-synuclein pre-formed fibrils^57,58^. While initially surprising, the reductions we found in MitoPark axons following stimulation may result from several mechanisms. First, it is possible that calcium channel expression is reduced in MitoPark axons relative to controls. This is consistent with a downregulation of other voltage-gated ion channels following axonal injury in other central and peripheral axons^59,60^. Furthermore, beyond overt downregulation, calcium channel organization may change, as well as the spatial relationship of calcium channels with other plasma membrane ion channels or intracellular calcium stores^61,62^. Calcium stores themselves may also be affected: impairments in calcium storage by the endoplasmic reticulum have been associated with PD^53^. Additionally, the disruption of mitochondrial quality control by *Tfam* deletion in our model mice should impair the ability of mitochondria to regulate intracellular calcium via reuptake^24,40^. However, we would expect disruption in any calcium storage or other buffering to result in both higher baseline calcium levels as well as slower decays of calcium signals. As we did not observe either of these in our photometry or two-photon data, it is unlikely that alterations in calcium storage or buffering drive the deficit we observed in stimulation-evoked calcium influx in MitoPark axons. Nevertheless, it is possible that calcium storage is impaired in MitoPark axons in ways that do not affect activity-dependent calcium dynamics but may influence other axonal functions, including trafficking and synaptic transmission^62,63^. An additional factor that may affect cytosolic calcium is the adenosine-triphospate-driven calcium pumps (Ca-ATPases), which largely serve to remove calcium from the intracellular space^64^. However, we would expect mitochondrial impairment in the MitoPark mice to lead to reduced ATP levels and reduced function of these Ca-ATPases, in turn increasing cytosolic calcium at baseline, which is opposite to the decrease in baseline calcium that we observed in these mice. In the absence of changes in buffering or ATP-driven changes in calcium pump function, the most likely candidate mechanisms for the reduction in calcium influx we see in MitoPark axons are a reduction in calcium channel density, a change in AP shape that leads to reduced per-AP calcium influx, or some combination of the two.

The advent of genetically-encoded calcium sensors and photometry techniques has vastly expanded our ability to understand the behavior of axons in both *in vivo* and *ex vivo* contexts. The small size of most central nervous system axons makes them relatively inaccessible to electrophysiological techniques that are used to examine excitability in cell bodies and dendrites. Here, we combine both photometry techniques, which measure the activity of many axons, with two-photon imaging and electrophysiology, which each sample one axon at a time. Each technique offers advantages and disadvantages, and by combining them to examine the same core question, it became apparent that the data collected with each technique can seem contradictory. We expect that the differences we see are a product of the slightly different focuses of each approach. When sampling across all axons in the field of view, as with photometry, even axons that are not responsive to the stimulation will contribute to the calcium signal, namely by raising the baseline calcium and reducing the signal -to-noise ratio. With two-photon imaging, we selected for only the axons that were responsive to stimulation, increasing our signal-to-noise but not accounting for axons that are not responsive to stimulation. Because nigrostriatal axons have branching architecture, it is not possible to differentiate between axons that are not responsive because the stimulating electrode is not positioned to stimulate them versus those axons that are not responsive due to excitability deficits. This same selection issue also applied for electrophysiological recordings, where a non-responsive axon could not be easily classified as stimulated and failing to propagate versus simply not stimulated. For example, the apparent failures in propagation at 2 Hz are more pronounced in photometry data than in our cell-attached recordings. It is possible that, by selecting for responsive axons, we overestimated the fidelity of AP propagation in MitoPark axons. Nevertheless, we believe that the multimodal approach we used here provides more information than photometry alone, and future studies should try, when possible, to gather data using both population-level and individual axon-level approaches.

In this study, we show evidence for AP propagation failures in nigrostriatal axons using cell-attached recordings of axonal varicosities. Cable theory predicts that propagation failures are likely to occur at branch points, and modeling studies have shown that increasing branch order is associated with higher propagation failure rates^8,65^. However, whether dopaminergic axons exhibit AP conduction failures has been a matter of debate in the field. For example, spiking activity in VTA dopamine cell bodies does not correlate with dopamine release in the nucleus accumbens in a reward-learning task^66^. Conversely, in molecularly-defined midbrain cell types, somatic firing did correlate with striatal axon calcium influx during behavior^67^. Recently, branch-site AP failures have been detected in nigrostriatal axons using two-photon calcium imaging approaches^68^. Here, we show electrophysiological evidence of frequency-dependent propagation failures in nigrostriatal axons that differs by disease state. Our results showing AP propagation failure prompts questions about how ion channel organization enables nigrostriatal axons to maintain high-fidelity propagation despite challenges to propagation, such as highly-branching morphology and burst firing, as well as how this organization is disrupted in the case of injury, degeneration, or energetic challenge. For example, in both peripheral and central nervous system axons, injury induces reorganization of axonal ion channels^59–61,69^. Understanding how axonal ion channel organization and propagation fidelity change in tandem is crucial to unlocking new approaches to address diseases that stem from axonal injury.

In Parkinson’s disease, a major hurdle has been the lack of disease-modifying treatments for the condition. While deep-brain stimulation and dopamine replacement therapy provide symptom relief, attempts to restore dopaminergic inputs to the striatum, or prevent their death altogether, have so far been largely unsuccessful. Because axons degenerate first, promoting their survival may be an avenue for early intervention. However, without knowledge of how axonal function changes before and during this degeneration, it is difficult to know what kinds of interventions have the highest chance of success. Our work in a mouse model of parkinsonism shows that the earliest deficits of axonal function in this model are deficits in calcium influx and excitability. Whether this is a common feature across multiple PD models and in PD itself remains to be explored, but this research provides a starting point for exploring new disease-modifying treatments of PD.

## ACKNOWLEDGMENTS

This research was supported by the Intramural Research Program (ZIANS003135) of the National Institutes of Health (NIH), National Institute of Neurological Disorders and Stroke (NINDS). The contributions of the NIH authors were made as part of their official duties as NIH federal employees, are in compliance with agency policy requirements, and are considered Works of the United States Government. However, the findings and conclusions presented in this paper are those of the authors and do not necessarily reflect the view of the NIH or the U.S. Department of Health and Human Services. The authors would like to thank Vincent Schram and Ling Yi from the NICHD Microscopy and Imaging Core for their assistance with slide scanner imaging experiments. They also thank members of the Khaliq lab at NINDS for helpful discussion and feedback.

## SUPPLEMENTAL MATERIALS

**Supplemental Figure 1:**
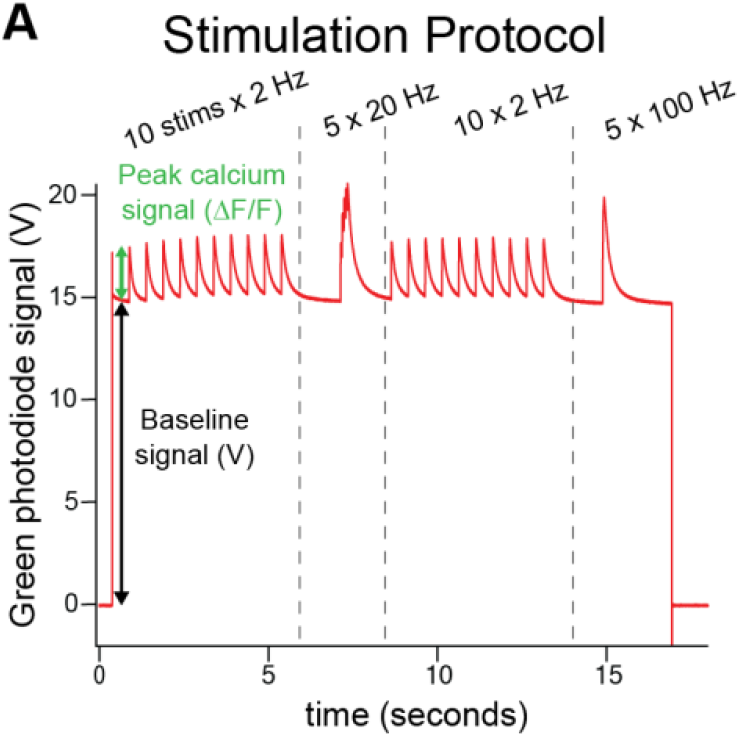
Nigrostriatal axons from MitoPark mice show an age-dependent reduction in baseline calcium signals. *A.* Stimulation protocol used to measured evoked calcium influx in nigrostriatal axons. Baseline calcium was measured as the photodiode signal, in volts, after the photodiode turned on but before activity was stimulated.

**Supplemental Figure 2:**
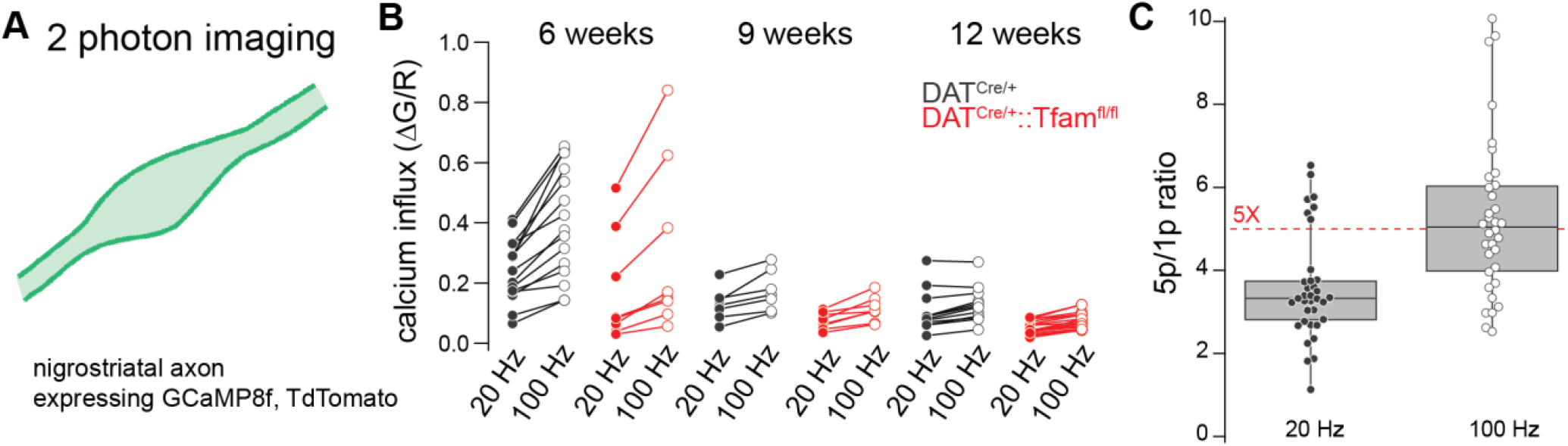
Calcium influx does not saturate at 20 Hz in nigrostriatal axons. ***A.*** Schematic showing experimental approach. Calcium influx was measured for single axons using two-photon laser microscopy. Axons were stimulated with 5 pulses at 20 Hz and 5 pulses at 100 Hz. ***B.*** Summary of calcium influx measured in single axons in response to trains of 20 and 100 Hz stimulation. Lines connect signals recorded from the same axon. In all ages and genotypes, 100 Hz stimulation leads to greater calcium influx than 20 Hz stimulation (p < 0.05, Wilcoxon matched-pairs signed rank tests). ***C.*** The ratio of peak calcium influx at the fifth pulse relative to the calcium associated with the first pulse shows that AP-associated calcium influx in single axons is largely within the linear range of the jGCaMP8f sensor.

**Supplemental Figure 3:**
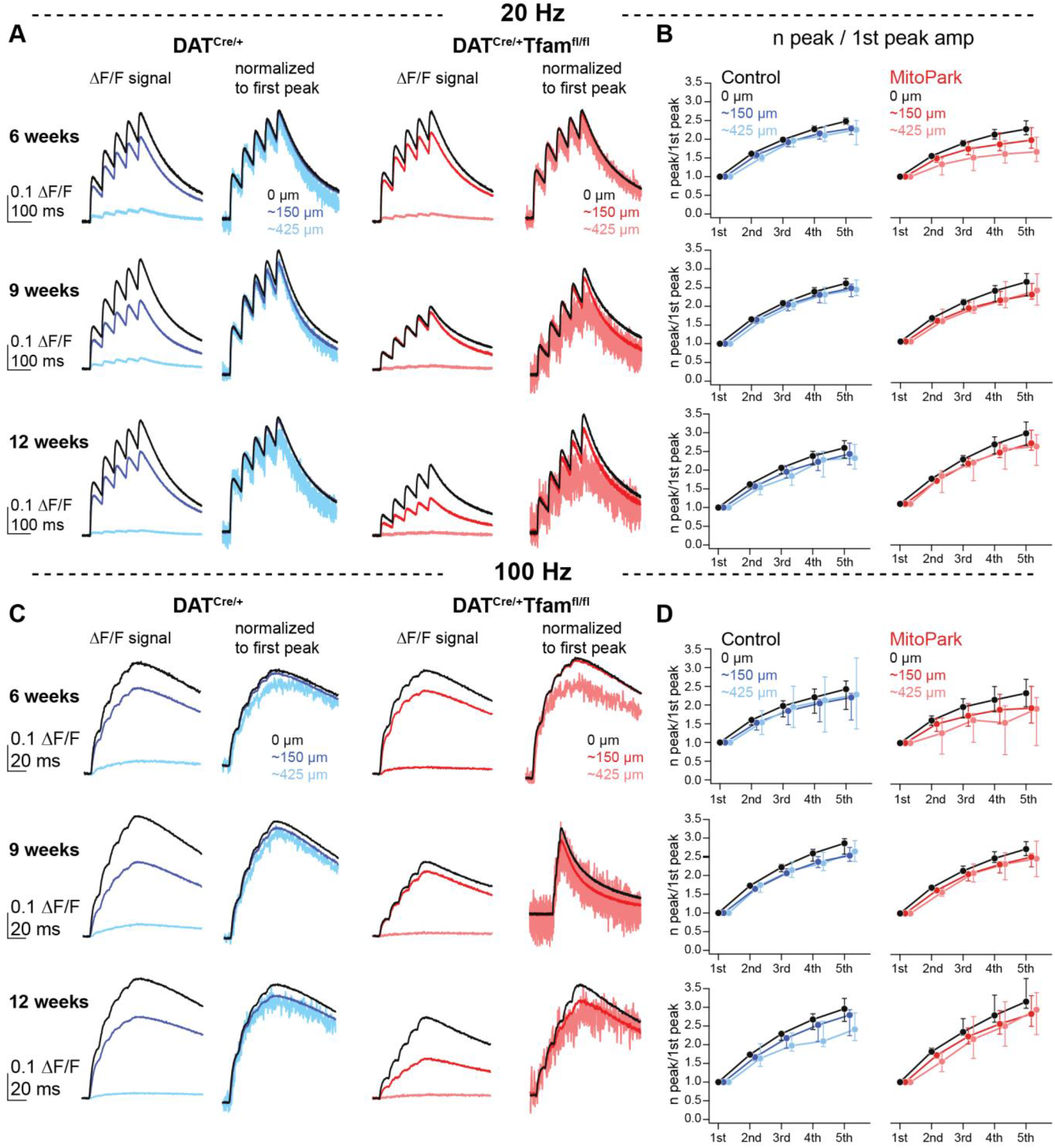
Calcium influx decreases with distance for high-frequency bursts in nigrostriatal axons. ***A.*** Left column: Representative ΔF/F signals for bursts (5 pulses) at 20 Hz in DAT^Cre/+^ (blues) and DAT^Cre/+^::Tfam^fl/fl^ (reds). Right column: Same traces shown in left column normalized to the first of five pulses. ***B.*** Relationship between peak number and normalized calcium influx for both genotypes at three distances, 0 µm from stimulating electrode (black), 150 µm (blue, red), and 425 µm (light blue, light red). Data are shown as median and interquartile range. ***C.*** Same as panel A but for 5 pulses of 100 Hz stimulation. ***D.*** Peak number and normalized calcium influx for 100 Hz stimulation at different distances from the stimulating electrode.

**Supplemental Figure 4:**
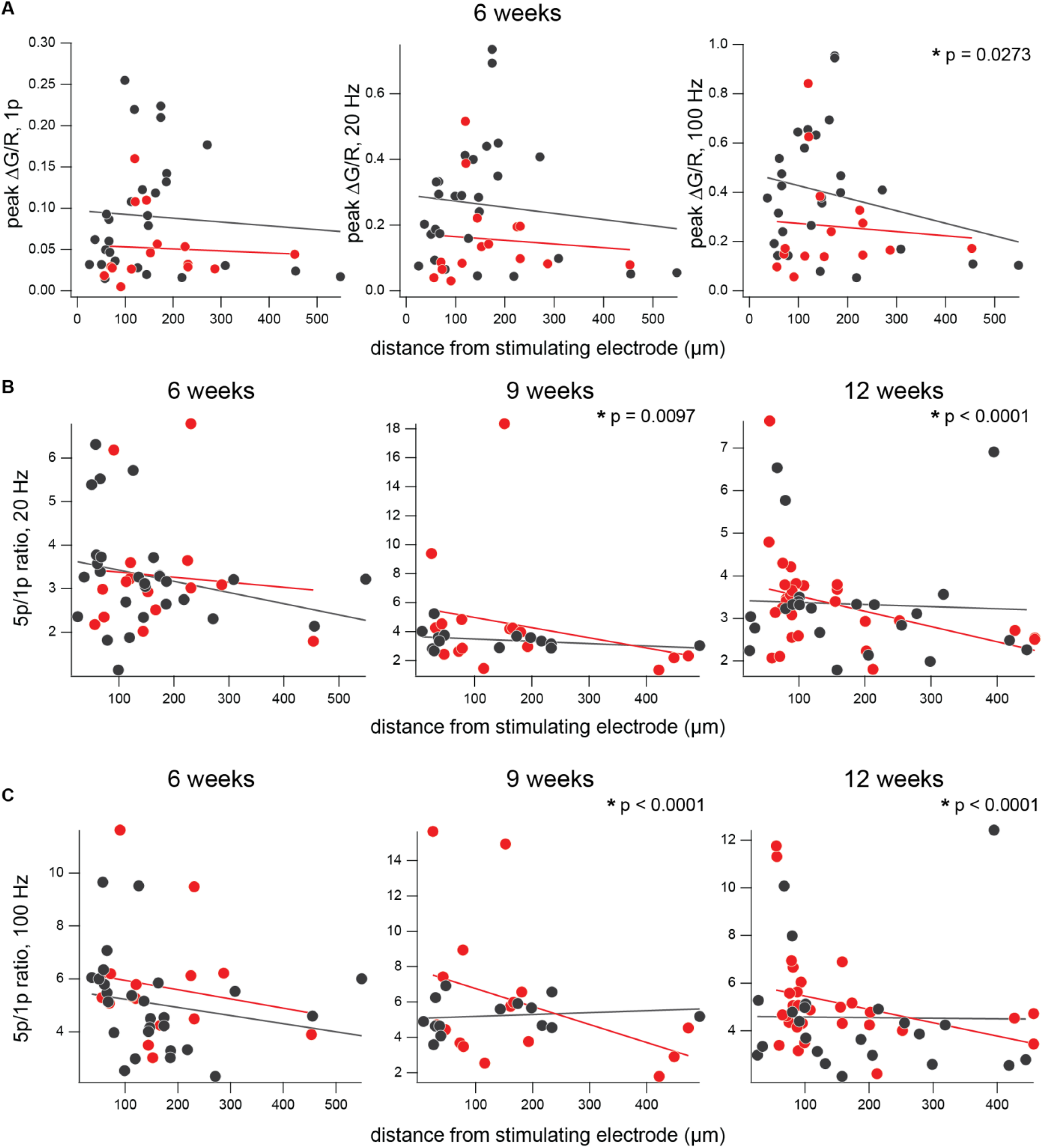
Calcium influx decreases in a frequency- and distance-dependent manner in MitoPark axons. ***A.*** Relationship between peak calcium influx and distance from the stimulating electrode for single pulses, 20 Hz, and 100 Hz stimulation in 6-week animals of both genotypes. The line of best fit was calculated for each genotype at each stimulation frequency. ***B.*** The relationship between 5p/1p ratio and distance for 20 Hz stimulation at each timepoint. At 9 and 12 weeks, MitoPark axons showed a distance-dependent decrease in this ratio. *C.* As in B, but for 100 Hz stimulation. As above, 9- and 12-week MitoPark axons showed a negative correlation between 5p/1p ratio and distance, while control axons did not.

**Supplemental Figure 5:**
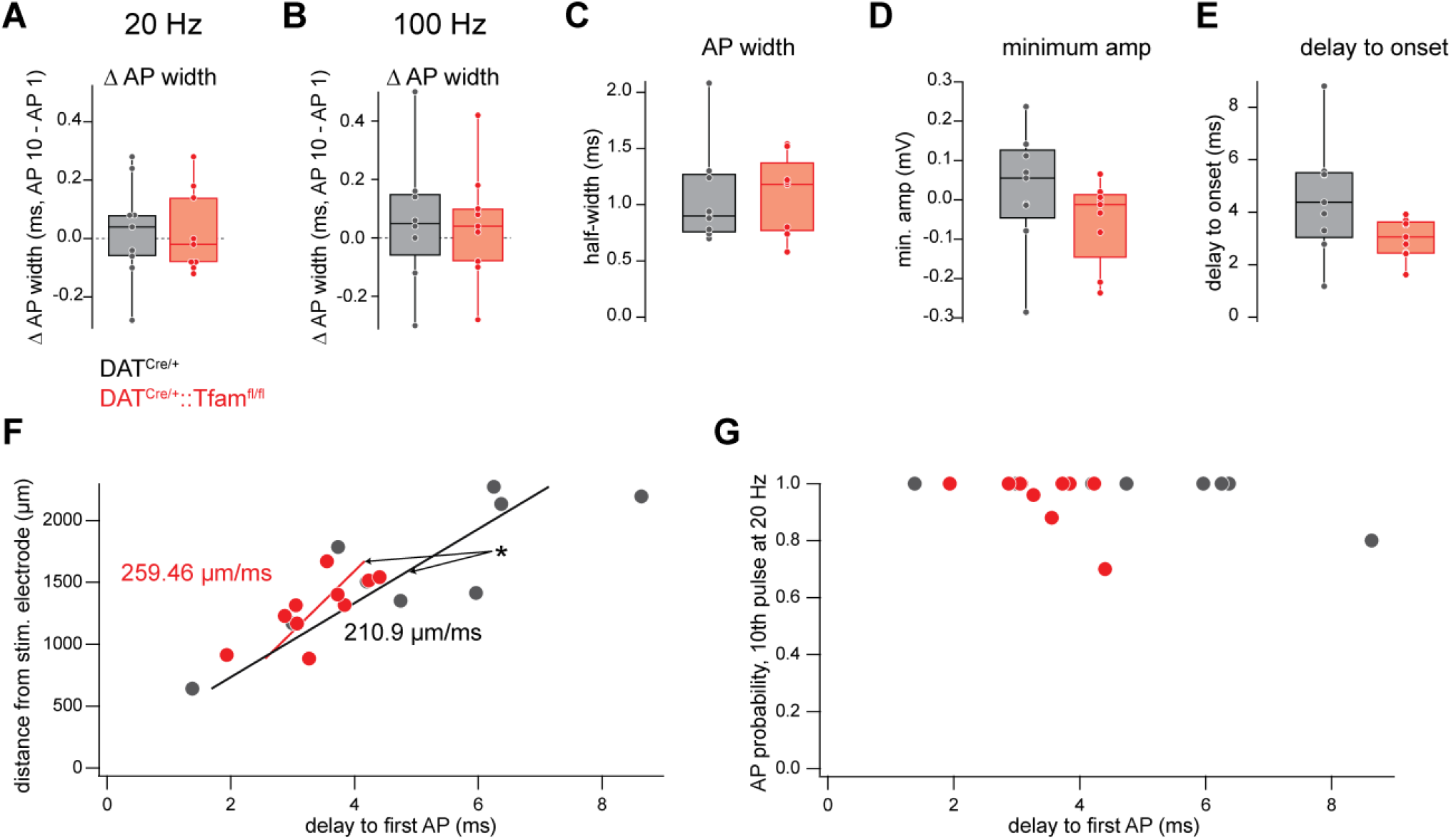
Cell-attached AP properties do not differ between genotype at 9 weeks. ***A.*** AP width, measured as the time between cell-attached AP maximum amplitude to minimum amplitude trough, did not change with 20 Hz stimulation. ***B.*** Both MitoPark and control axons show no change in AP width with 100 Hz firing. ***C.*** Overall, there was no difference in cell-attached AP width between MitoPark and control axons. ***D.*** The minimum amplitude of the cell-attached AP waveform across genotypes at 9 weeks. ***E.*** Delay to AP onset, measured from stimulus offset to AP maximum amplitude. ***F.*** The relationship between delay to AP onset and distance from the stimulation electrode showed a positive correlation in both genotypes. ***G.*** The relationship between delay to AP onset and AP probability for the 10^th^ AP in a 20 Hz burst for both genotypes.

## REFERENCES

1. Haddad, D. & Nakamura, K. Understanding the susceptibility of dopamine neurons to mitochondrial stressors in Parkinson’s disease. FEBS Lett. 589, 3702–3713 (2015).

2. Gonzalez-Rodriguez, P., Zampese, E. & Surmeier, D. J. Selective neuronal vulnerability in Parkinson’s disease. Prog. Brain Res. 252, 61–89 (2020).

3. Sulzer, D. & Surmeier, D. J. Neuronal vulnerability, pathogenesis, and Parkinson’s disease. Movement Disorders 28, 715–724 (2013).

4. Bolam, J. P. & Pissadaki, E. K. Living on the edge with too many mouths to feed: Why dopamine neurons die. Movement Disorders 27, 1478–1483 (2012).

5. Surmeier, D. J., Obeso, J. A. & Halliday, G. M. Selective neuronal vulnerability in Parkinson disease. Nature Reviews Neuroscience 2017 18:2 18, 101–113 (2017).

6. Matsuda, W. et al. Single Nigrostriatal Dopaminergic Neurons Form Widely Spread and Highly Dense Axonal Arborizations in the Neostriatum. Journal of Neuroscience 29, 444–453 (2009).

7. Grace, A. A. & Bunney, B. S. The control of firing pattern in nigral dopamine neurons: single spike firing. Journal of Neuroscience 4, 2866–2876 (1984).

8. Pissadaki, E. K. & Bolam, J. P. The energy cost of action potential propagation in dopamine neurons: clues to susceptibility in Parkinson’s disease. 10.3389/fncom.2013.00013 (2013) doi:10.3389/fncom.2013.00013.

9. Braak, H., Ghebremedhin, E., Rüb, U., Bratzke, H. & Del Tredici, K. Stages in the development of Parkinson’s disease-related pathology. Cell and Tissue Research 2004 318:1 318, 121–134 (2004).

10. Attwell, D. & Laughlin, S. B. An energy budget for signaling in the grey matter of the brain. Journal of Cerebral Blood Flow and Metabolism 21, 1133–1145 (2001).

11. Howarth, C., Gleeson, P. & Attwell, D. Updated energy budgets for neural computation in the neocortex and cerebellum. Journal of Cerebral Blood Flow and Metabolism 32, 1222–1232 (2012).

12. Hasenstaub, A., Otte, S., Callaway, E. & Sejnowski, T. J. Metabolic cost as a unifying principle governing neuronal biophysics. Proc. Natl. Acad. Sci. U. S. A. 107, 12329–12334 (2010).

13. Pacelli, C. et al. Elevated Mitochondrial Bioenergetics and Axonal Arborization Size Are Key Contributors to the Vulnerability of Dopamine Neurons. Current Biology 25, 2349–2360 (2015).

14. Angelova, P. R. & Abramov, A. Y. Role of mitochondrial ROS in the brain: from physiology to neurodegeneration. FEBS Lett. 592, 692–702 (2018).

15. Desai, V. G., Feuers, R. J., Hart, R. W. & Ali, S. F. MPP+-induced neurotoxicity in mouse is age-dependent: Evidenced by the selective inhibition of complexes of electron transport. Brain Res. 715, 1–8 (1996).

16. Nicklas, W. m. dot J., Vyas, I. & Heikkila, R. E. Inhibition of NADH-linked oxidation in brain mitochondria by 1-methyl-4-phenyl-pyridine, a metabolite of the neurotoxin, 1-methyl-4-phenyl-1,2,5,6-tetrahydropyridine. Life Sci. 36, 2503–2508 (1985).

17. Betarbet, R. et al. Chronic systemic pesticide exposure reproduces features of Parkinson’s disease. Nat. Neurosci. 3, 1301–1306 (2000).

18. Schapira, A. H. V. et al. Mitochondrial Complex I Deficiency in Parkinson’s disease. The Lancet 333, 1269 (1989).

19. Mann, V. M. et al. Brain, skeletal muscle and platelet homogenate mitochondrial function in Parkinson’s disease. Brain 115 **(Pt** **2**), 333–342 (1992).

20. Janetzky, B. et al. Unaltered aconitase activity, but decreased complex I activity in substantia nigra pars compacta of patients with Parkinson’s disease. Neurosci. Lett. 169, 126–128 (1994).

21. Henrich, M. T., Oertel, W. H., Surmeier, D. J. & Geibl, F. F. Mitochondrial dysfunction in Parkinson’s disease – a key disease hallmark with therapeutic potential. Molecular Neurodegeneration 2023 18:1 18, 83-(2023).

22. Narendra, D. P. & Youle, R. J. The role of PINK1–Parkin in mitochondrial quality control. Nature Cell Biology 2024 26:10 26, 1639–1651 (2024).

23. González-Rodríguez, P. et al. Disruption of mitochondrial complex I induces progressive parkinsonism. Nature 2021 599:7886 599, 650–656 (2021).

24. Ekstrand, M. I. et al. Progressive parkinsonism in mice with respiratory-chain-deficient dopamine neurons. Proceedings of the National Academy of Sciences 104, 1325–1330 (2007).

25. Burke, R. E. & O’Malley, K. Axon Degeneration in Parkinson’s Disease. Exp. Neurol. 246, 72 (2013).

26. Hornykiewicz, O. Biochemical aspects of Parkinson’s disease. Neurology 51, S2–S9 (1998).

27. Kordower, J. H. et al. Disease duration and the integrity of the nigrostriatal system in Parkinson’s disease. Brain 136, 2419–2431 (2013).

28. O’Malley, K. L. The Role of Axonopathy in Parkinson’s Disease. Exp. Neurobiol. 19, 115 (2010).

29. Kurowska, Z. et al. Is Axonal Degeneration a Key Early Event in Parkinson’s Disease? J. Parkinsons Dis. 6, 703–707 (2016).

30. Cooper, O. et al. Pharmacological rescue of mitochondrial deficits in iPSC-derived neural cells from patients with familial Parkinson’s disease. Sci. Transl. Med. 4, (2012).

31. Mallach, A. et al. Post mortem examination of Parkinson’s disease brains suggests decline in mitochondrial biomass, reversed by deep brain stimulation of subthalamic nucleus. FASEB J. 33, 6957– 6961 (2019).

32. Larsson, N. G. et al. Mitochondrial transcription factor A is necessary for mtDNA maintenance and embryogenesis in mice. Nat. Genet. 18, 231–236 (1998).

33. Ekstrand, M. I. & Galter, D. The MitoPark Mouse – An animal model of Parkinson’s disease with impaired respiratory chain function in dopamine neurons. Parkinsonism Relat. Disord. 15, S185–S188 (2009).

34. Fushiki, A., et al. A Vulnerable Subtype of Dopaminergic Neurons Drives Early Motor Deficits in Parkinson’s Disease. bioRxiv 2024.12.20.629776 (2026) doi:10.1101/2024.12.20.629776.

35. Good, C. H. et al. Impaired nigrostriatal function precedes behavioral deficits in a genetic mitochondrial model of Parkinson’s disease. The FASEB Journal 25, 1333–1344 (2011).

36. Branch, S. Y. et al. Dopaminergic Neurons Exhibit an Age-Dependent Decline in Electrophysiological Parameters in the MitoPark Mouse Model of Parkinson’s Disease. Journal of Neuroscience 36, 4026– 4037 (2016).

37. O’Neill, B., Patel, J. C. & Rice, M. E. Characterization of Optically and Electrically Evoked Dopamine Release in Striatal Slices from Digenic Knock-in Mice with DAT-Driven Expression of Channelrhodopsin. ACS Chem. Neurosci. 8, 310–319 (2017).

38. Kramer, P. F. et al. Synaptic-like axo-axonal transmission from striatal cholinergic interneurons onto dopaminergic fibers. Neuron 110, 2949–2960 (2022).

39. Rademacher, K. et al. Chronic hyperactivation of midbrain dopamine neurons causes preferential dopamine neuron degeneration. Elife 13, (2025).

40. Beckstead, M. J. & Howell, R. D. Progressive parkinsonism due to mitochondrial impairment: Lessons from the MitoPark mouse model. Exp. Neurol. 341, 113707 (2021).

41. Grace, A. A. & Bunney, B. S. The control of firing pattern in nigral dopamine neurons: burst firing. Journal of Neuroscience 4, 2877–2890 (1984).

42. Paladini, C. A. & Roeper, J. Generating bursts (and pauses) in the dopamine midbrain neurons. Neuroscience 282, 109–121 (2014).

43. Attwell, D. & Gibb, A. Neuroenergetics and the kinetic design of excitatory synapses. Nature Reviews Neuroscience 2005 6:11 6, 841–849 (2005).

44. Yi, G., Wang, J., Wei, X. & Che, Y. Energy cost of action potential generation and propagation in thalamocortical relay neurons during deep brain stimulation. IEEE Trans. Biomed. Eng. 66, 3457–3471 (2019).

45. Qian, K., Yu, N., Tucker, K. R., Levitan, E. S. & Canavier, C. C. Mathematical analysis of depolarization block mediated by slow inactivation of fast sodium channels in midbrain dopamine neurons. J. Neurophysiol. 112, 2779–2790 (2014).

46. Tucker, K. R., Huertas, M. A., Horn, J. P., Canavier, C. C. & Levitan, E. S. Pacemaker rate and depolarization block in nigral dopamine neurons: a somatic sodium channel balancing act. J. Neurosci. 32, 14519–14531 (2012).

47. Barcomb, K. et al. Overexpression of α–synuclein in Midbrain Dopamine Neurons Reduces Dopamine Release Without Cell Loss and Drives Mild Motor Deficits in Mice. bioRxiv 2025.03.17.643840 (2025) doi:10.1101/2025.03.17.643840.

48. Dovonou, A. et al. Animal models of Parkinson’s disease: bridging the gap between disease hallmarks and research questions. Translational Neurodegeneration 2023 12:1 12, 36-(2023).

49. Yue, M. et al. Progressive dopaminergic alterations and mitochondrial abnormalities in LRRK2 G2019S knock-in mice. Neurobiol. Dis. 78, 172–195 (2015).

50. Liu, C. & Kaeser, P. S. Mechanisms and regulation of dopamine release. Curr. Opin. Neurobiol. 57, 46 (2019).

51. Liu, C., Kershberg, L., Wang, J., Schneeberger, S. & Kaeser, P. S. Dopamine Secretion Is Mediated by Sparse Active Zone-like Release Sites. Cell 172, 706–718.e15 (2018).

52. Lebowitz, J. J. et al. Synaptotagmin-1 is a Ca2+ sensor for somatodendritic dopamine release. Cell Rep. 42, 111915 (2023).

53. Wolfart, J., Neuhoff, H., Franz, O. & Roeper, J. Differential Expression of the Small-Conductance, Calcium-Activated Potassium Channel SK3 Is Critical for Pacemaker Control in Dopaminergic Midbrain Neurons. Journal of Neuroscience 21, 3443–3456 (2001).

54. Zaichick, S. V., McGrath, K. M. & Caraveo, G. The role of Ca2+ signaling in Parkinson’s disease. Dis. Model. Mech. 10, 519–535 (2017).

55. Ricke, K. M. et al. Mitochondrial Dysfunction Combined with High Calcium Load Leads to Impaired Antioxidant Defense Underlying the Selective Loss of Nigral Dopaminergic Neurons. Journal of Neuroscience 40, 1975–1986 (2020).

56. Puopolo, M., Raviola, E. & Bean, B. P. Roles of Subthreshold Calcium Current and Sodium Current in Spontaneous Firing of Mouse Midbrain Dopamine Neurons. Journal of Neuroscience 27, 645–656 (2007).

57. Wilson, C. J. & Callaway, J. C. Coupled oscillator model of the dopaminergic neuron of the substantia nigra. J. Neurophysiol. 83, 3084–3100 (2000).

58. Dryanovski, D. I. et al. Calcium Entry and α-Synuclein Inclusions Elevate Dendritic Mitochondrial Oxidant Stress in Dopaminergic Neurons. Journal of Neuroscience 33, 10154–10164 (2013).

59. Chan, C. S. et al. ‘Rejuvenation’ protects neurons in mouse models of Parkinson’s disease. Nature 2007 447:7148 447, 1081–1086 (2007).

60. Mert, T. Roles of axonal voltage-dependent ion channels in damaged peripheral nerves. Eur. J. Pharmacol. 568, 25–30 (2007).

61. Rizzo, M. A., Kocsis, J. D. & Waxman, S. G. Selective loss of slow and enhancement of fast Na+currents in cutaneous afferent dorsal root ganglion neurones following axotomy. Neurobiol. Dis. 2, 87–96 (1995).

62. Romer, S. H. et al. Redistribution of Kv2.1 ion channels on spinal motoneurons following peripheral nerve injury. Brain Res. 1547, 1–15 (2014).

63. Panzera, L. C. et al. Activity-dependent endoplasmic reticulum Ca2+ uptake depends on Kv2.1-mediated endoplasmic reticulum/plasma membrane junctions to promote synaptic transmission. Proc. Natl. Acad. Sci. U. S. A. 119, e2117135119 (2022).

64. Kole, K. & Kole, M. H. P. Robust activity-dependent mitochondrial calcium dynamics at the AIS is dispensable for action potential generation. J. Physiol. 604, 3032–3052 (2026).

65. Zampese, E. & Surmeier, D. J. Calcium, Bioenergetics, and Parkinson’s Disease. Cells 2020, *Vol. 9, Page* 2045 9, 2045 (2020).

66. Debanne, D. Information processing in the axon. Nature Reviews Neuroscience 2004 5:4 5, 304–316 (2004).

67. Mohebi, A. et al. Dissociable dopamine dynamics for learning and motivation. Nature 2019 570:7759 570, 65–70 (2019).

68. Azcorra, M. et al. Unique functional responses differentially map onto genetic subtypes of dopamine neurons. Nature Neuroscience 2023 1–13 (2023) doi:10.1038/s41593-023-01401-9.

69. Yee, A. G., Liao, Y., Muntean, B. S. & Ford, C. P. Discrete spatiotemporal encoding of striatal dopamine transmission. Science (1979). 389, 200–206 (2025).

70. Henry, M. A., Freking, A. R., Johnson, L. R. & Levinson, S. R. Sodium channel Nav1.6 accumulates at the site of infraorbital nerve injury. BMC Neuroscience 2007 8:1 8, 56-(2007).

